# METTL3/MYCN cooperation drives neural crest differentiation and provides therapeutic vulnerability in neuroblastoma

**DOI:** 10.1101/2023.10.06.561194

**Authors:** Ketan Thombare, Roshan Vaid, Perla Pucci, Akram Mendez, Rebeca Burgos-Panadero, Ritish Ayyalusamy, Aqsa Ali Rehan, Mohammad Hassan Baig, Sagar Dattatraya Nale, Christoph Bartenhagen, Jae-June Dong, Matthias Fischer, Suzanne D. Turner, Tanmoy Mondal

## Abstract

Neuroblastoma (NB) is the most common extracranial childhood cancer, caused by the improper differentiation of developing trunk neural crest cells (tNCC) in the sympathetic nervous system. The *N*^6^-methyladenosine (m^6^A) epitranscriptomic modification controls post-transcriptional gene expression but the mechanism by which the m^6^A methyltransferase complex METTL3/METTL14/WTAP is recruited to specific loci remains to be fully characterized. We explored whether the m^6^A epitranscriptome could fine-tune gene regulation in migrating/differentiating tNCC. We demonstrate that the m^6^A modification regulates the expression of *HOX* genes in tNCC, thereby contributing to their timely differentiation into sympathetic neurons. Furthermore, we show that posterior *HOX* genes are m^6^A modified in MYCN-amplified NB with reduced expression. In addition, we provide evidence that sustained overexpression of the MYCN oncogene in tNCC drives METTL3 recruitment to a specific subset of genes including posterior *HOX* genes creating an undifferentiated state. Moreover, METTL3 depletion/inhibition induces DNA damage and differentiation of MYCN overexpressing cells and increases vulnerability to chemotherapeutic drugs in MYCN-amplified patient-derived xenografts (PDX) cells, suggesting METTL3 inhibition could be a potential therapeutic approach for NB.

## Introduction

RNA modification (also known as epitranscriptomics) can control important steps in RNA biogenesis such as RNA stability and RNA transport. One of the most abundant modifications of cellular RNA is *N^6^*-methyladenosine (m^6^A). m^6^A is deposited on cellular RNA co-transcriptionally by the enzyme complex METTL3/METTL14/WTAP (1). A role for m^6^A RNA modification has been shown in chromatin regulation, stress response, DNA damage repair, as well as in viral infection by regulating gene expression both at transcriptional and post-transcriptional levels (2–5). Active chromatin modification H3K36me3, RNA binding proteins such as RBFOX2, and transcription factors have been implicated in METTL3 recruitment (6–8). Despite these studies, the mechanism which drives the locus specific recruitment of METTL3 remains unclear.

Neuroblastoma (NB) is a heterogeneous disease, it can spontaneously regress but in cases of high-risk disease, even with intensive therapy, disease relapse is not uncommon. As such, a better understanding of this disease and novel therapeutic strategies are urgently required. Amplification of the MYCN oncogene is one of the major genetic alterations found in NB and correlates with poor survival (9,10). MYCN amplification creates an undifferentiated state in NB (10,11), but detailed molecular mechanisms are lacking particularly as to how deregulation of MYCN creates an undifferentiated state in early developing human trunk neural crest cells (tNCC). MYCN overexpression has been shown to induce the transformation of neural crest cells into NB cells in humanized mouse models (12–14). Recently tNCC were derived from human embryonic stem cells (hESC) *in vitro* and these tNCC can be driven further to sympathoadrenal progenitors (SAP) and sympathetic neurons (SN) (15,16). Whether m^6^A modification has any role in this differentiation process has not been investigated.

METTL3 has been shown to have a tumor promoting role in many cancers (17) and its inhibition through small molecules has recently been proposed as a therapeutic strategy for acute myeloid leukemia (AML) (18). A role for METTL3 mediated m^6^A modification was recently reported in Alternative Lengthening of Telomeres-positive (ALT+) NB (19) but its function is unknown in other types of high-risk NB. High expression levels of METTL3 are predictive of an inferior outcome for NB patients and METTL3 is expressed in both ALT+ and high-risk MYCN-amplified NB tumors suggesting it may have broader relevance (19).

In this study we showed higher expression levels of METTL3 in tNCC compared to hESC, correlating with an increase in overall m^6^A peaks in tNCC. We also showed that METTL3 regulates the timely differentiation of tNCC by regulating *HOX* gene expression. We observed that MYCN overexpression can lead to an undifferentiated state in tNCC by downregulating posterior *HOX* gene expression. This MYCN mediated undifferentiated state can be reversed by METTL3 depletion suggesting that METTL3 inhibition could be a novel therapeutic option for high-risk NB.

## Materials and Methods

### Neuroblastoma (NB) cell lines and culture conditions

MYCN-amplified NB cell lines SK-N-BE(2), IMR-32, and NGP were used in this study. SK-N-BE(2) and NGP were procured from DSMZ whereas IMR-32 from CLS Cell Lines Service. SHEP cells were a gift from Dr. Marie Arsenian-Henriksson (Karolinska Institute, Sweden). SK-N-BE(2) cells were cultured in DMEM/F-12 media supplemented with 10% FBS, 1x GlutaMAX, and penicillin/streptomycin. NGP and SHEP cells were cultured in Dulbecco’s modified Eagle’s medium (DMEM) supplemented with 10% fetal bovine serum (FBS) and penicillin/streptomycin. IMR-32 cells were cultured in Minimum Essential Medium (MEM) supplemented with 10% FBS, 1 mM sodium pyruvate, 1x GlutaMAX (Gibco), and penicillin/streptomycin. Patient-derived xenografts (PDX) cells, COG-N-496h (MYCN-amplified, ALK WT, P53 WT) were obtained from the Children’s Oncology Group (Texas, USA) and cultured in Iscove’s Modified Dulbecco’s Medium (IMDM) plus 20% FBS, 4mM L-Glutamine, 1X ITS (5 µg/mL insulin, 5 µg/mL transferrin, 5 ng/mL selenous acid). Cells were confirmed to be mycoplasma-negative using the MycoAlert Mycoplasma detection assay (Lonza, LT07218) and maintained in a humidified incubator at 37°C with 5% CO_2_.

### hESC culture and differentiation

Human embryonic stem cell line WA09 (H9) obtained from Dr. Fredrik H. Sterky (Sahlgrenska University Hospital, Gothenburg, Sweden) were cultured on Matrigel-coated plates with iPS-Brew XF (Miltenyi) media. To differentiate hESC to trunk neural crest cells (tNCC), hESC were dissociated using Accutase and seeded (day 0) on Matrigel-coated plates to induce differentiation to neuromesodermal progenitor cells (NMP, day 3) which were then driven to tNCC (days 7-9), sympathoadrenal progenitors (SAP, day 12), and further towards sympathetic neurons (SN, day 19-25) as described previously (15,16). For neural crest stem cells (NCSC) induction, hESCs were dissociated using Accutase and seeded (day 0) on Matrigel-coated plates at a density of 2×10^4^ cells/cm^2^ in iPS-Brew XF media with ROCK inhibitor (Y-27632 10 μM). Following day media was replaced with NCSC differentiation medium as previously described (20). NCSC differentiation medium was replaced every 2 days and cells were passaged on Matrigel-coated plates every 3-4 days of reaching 90% confluency. Reagents used for hESC maintenance and differentiation are listed in Supplementary Table S1.

### Transient gene silencing and overexpression

METTL3 siRNAs were used to induce transient knockdown in SK-N-BE(2) cells with RNAiMAX (ThermoFisher) reagent, following the manufacturer’s guidelines. HOXC8 and HOXC9 plasmids were transfected using Lipofectamine 3000 reagent (ThermoFisher), following the manufacturer’s guidelines. Target siRNA sequences and plasmid information are provided in Supplementary Table S1.

### METTL3 shRNA lentiviral packaging and viral transduction

shRNA constructs targeting METTL3 and control shRNA in the pLKO.1 vector were procured from Sigma. To generate lentiviral particles for each shMETTL3 and control shRNA construct, HEK293T cells were co-transfected with these plasmids, along with pMD2.G and psPAX2 packaging plasmids, employing the CalPhos mammalian transfection kit (Takara Bio). Supernatants were harvested at 48 and 72 h post-transfection and subsequently stored at −80°C.

The METTL3 shRNA sequences were subsequently cloned into a tet-pLKO-puro vector and packaged to produce doxycycline (Dox)-inducible viral particles. Before selection with 1 µg/ml puromycin, NB cells, and hESC were transduced with shMETTL3 or shCtrl viral particles for 48 h. For inducible METTL3 KD, Dox was administered at a concentration of 2 µg/ml during hESC differentiation and 200 ng/ml for NB cells. Detailed plasmid information is provided in Supplementary Table S1.

### MYCN overexpression

PB-TRE3G-MYCN and XLone-GFP (21) were acquired from Addgene. Plasmid information is provided in Supplementary Table S1. PB-TRE3G-MYCN was used to amplify MYCN sequence with 1x Flag-tag followed by cloning into XLone vector replacing GFP to get Dox inducible Flag-MYCN. SHEP cells were co-transfected (1:1) with Flag-MYCN and pCYL43 piggyBac transposase using Lipofectamine 3000 as per the manufacturer’s protocol. Stably transfected cells were selected by treatment with 5 µg/ml Blasticidin. hESC H9 were nucleofected with two plasmids (1:1) Flag-MYCN and pCYL43 piggyBac transposase to generate stable cells. Amaxa 4D-Nucleofector system (Lonza) was employed for nucleofection as per the manufacturer’s instructions. Nucleofected cells were selected for a week using 2.5 µg/ml Blasticidin. Cells were then sparsely seeded, and colonies were picked, expanded, and screened for MYCN expression. Overexpression of MYCN was confirmed on the protein level using immunoblotting after 48 h Dox induction. The clone which showed consistently high MYCN expression was used in further experiments. Dox was used at a concentration of 2 µg/ml during hESC differentiation and 200 ng/ml in NB cells for MYCN overexpression. Flag-MYCN cells were differentiated following the tNCC differentiation protocol as described above with or without Dox treatment. DMSO or METTL3 inhibitor STM2457 (10µM) was introduced during the differentiation protocol as described in the figure legends.

### Retinoic acid (RA) mediated differentiation of NB cells

SK-N-BE(2) or NGP cells were seeded at a density of 1×10^5^ or 2×10^5^ per well, respectively. The following day, cells were pre-treated with DMSO or STM2457 (10 µM) for 24 h and followed by RA (10 µM) for an additional 3 days. A similar procedure was followed in Dox inducible METTL3 KD cells, where cells were induced with Dox for 24 h before the addition of RA for the following 3 days.

### Immunofluorescence staining (IF)

Cells were fixed in 4% formaldehyde for 10 min at room temperature (RT) and then rinsed twice with PBS. Following this, cells were permeabilized with 0.25% Triton X-100 in PBS for 10 min, followed by two washes with 0.1% Tween 20 in PBS (PBST). Subsequently, cells were blocked for 1 h at RT in 3% BSA in PBST. Primary antibodies, including METTL3 (1:300), phospho-Histone H2A.X (Ser139) (1:500), phospho-RPA32 (Ser33) (1:300), peripherin (PRPH) (1:100), PHOX2B (1:100), Oct-3/4 (1:100), HOXC9 (1:100), anti-HOXC8 (1:300), FLAG (1:500), MYCN (1:1000), and beta-Tubulin Isotype III (TUBB3)(1:500), were applied and incubated overnight at 4°C. After primary antibody exposure, cells were washed three times for 5 min each with PBST and then incubated for 1 h in the dark at RT with secondary antibodies labeled with Alexa Fluor 488 and Alexa Fluor 555 fluorochromes (1:800). Following secondary antibody incubation, cells were washed again three times for 5 min each with PBST. Prolong Gold with DAPI (ThermoFisher) was added to each coverslip, mounted on a slide, and air-dried in the dark for nuclei detection. Slides were imaged using a fluorescence microscope EVOS FL Auto (ThermoFisher), and image analysis was performed with ImageJ. Antibody details are provided in Supplementary Table S1.

### Immunoblotting and Co-IP

Cells were lysed in RIPA buffer (ThermoFisher) with a protease inhibitor cocktail (ThermoFisher). Protein concentrations were determined by bicinchoninic acid assay (ThermoFisher). Equal protein amounts were loaded onto a 4-12% Bis-Tris gel (ThermoFisher) and transferred to a nitrocellulose membrane with a Trans-Blot Turbo system (BioRad). Membranes were blocked for 1 h in 5% non-fat dry milk in PBST and probed overnight with primary antibodies diluted in 5% blocking solution: METTL3 (1:200), METTL14 (1:2000), MYCN (1:1000), Vinculin (1:5000), GAPDH (1:5000), A-tubulin (1:5000). Membranes were then incubated with HRP-linked secondary antibodies (Cell Signaling) for 1 h, and signals were detected using SuperSignal West Pico PLUS Chemiluminescent Substrate (ThermoFisher). Blots were developed and quantified using a ChemiDoc system and ImageLab software (BioRad). Co-IP was performed with METTL3 and MYCN antibodies as described before (22). Antibody details are provided in Supplementary Table S1.

### Protein Stability Assay

To inhibit protein synthesis, we employed the protein synthesis inhibitor cycloheximide (CHX) at a concentration of 50 µg/ml. Cells were collected at the specified time points, and protein was subsequently extracted. The METTL3 protein level was quantified through immunoblotting.

### Proximity ligation assay (PLA)

PLA was performed in tNCC or SHEP*^MYCN^* cells after 24 h Dox induced overexpression of MYCN using a Duolink PLA kit (Sigma, DUO92014) according to the manufacturer’s protocol. As a background control, a single antibody was used in this assay. Briefly, cells were fixed for 10 min in 4% PFA at RT before being blocked with a blocking solution. The cells were treated with primary antibodies targeting METTL3 and MYCN or HOXC8 and HOXC9 for 1 h at 37°C, followed by incubation with PLA probes for 1 h at 37°C in a humidified chamber. After three washes, a ligation-ligase solution was added and incubated for 30 min at 37°C. The slides were incubated for 100 min in an amplification solution containing polymerase at 37°C in the dark. Finally, the cells were stained with Prolong Gold containing DAPI, and coverslips were mounted on a slide and air-dried. Fluorescence microscopy was used to capture the fluorescence images.

### Proliferation assay

In total, 1×10^4^ cells per well (for SK-N-BE(2), 5×10^3^ cells per well) were seeded on a 96-well plate to assess cellular proliferation. SK-N-BE(2), IMR-32, and SHEP*^MYCN^* Dox inducible shCtrl or shMETTL3 cells were seeded and induced on the following day by adding Dox 200 ng/ml up to 6 days. For the combination experiment in SK-N-BE(2) treatment with STM2457 (10 µM) or with doxorubicin was performed at indicated concentrations for 72 h. CellTiter 96 non-radioactive cell proliferation assay kit (Promega, G4000) was used to determine cell growth and the manufacturer’s instructions were followed. Absorbance was measured using a microplate reader Infinite 50 (Tecan, Austria).

For dose-response matrices, cells were treated with log-scale concentrations of STM2457 in addition to log-scale concentrations of doxorubicin, and the DMSO concentration was maintained at < 0.2%. The potential synergy between STM2457 and doxorubicin was evaluated by calculating the synergy score based on the Loewe model (23) using the synergy finder web application, https://synergyfinder.fimm.fi (24). The synergy score is calculated for each combination of drug concentrations and also as an overall value and is defined as: >10 = synergistic; Between −10 and 10 = additive; <-10 = antagonistic.

### Colony formation assay

In total, 1×10^3^ SK-N-BE(2) shCtrl and shMETTL3 cells were plated in a 6-well plate. Dox induction was started after 24 h and Dox media was replaced every 2-3 days up to 14 days. Cells were then fixed with 10% formaldehyde for 10 min and then stained with 0.2% crystal violet solution for 1 h at RT. Excess crystal violet solution was carefully washed, and the plate was allowed to air dry and visualized using ChemiDoc.

### RNA sequencing (RNA-seq) and m^6^A RNA Immunoprecipitation sequencing (m^6^A RIP-seq)

RNA was isolated from cells using TRIZOL reagent (ThermoFisher) and Direct-zol RNA Miniprep (ZYMO research). NB cell line, hESC, NCC, and tNCC RNA (15 µg) were spiked-in with Bacterial RNA (10 ng) before fragmentation using RNA Fragmentation Reagents (ThermoFisher). For NB tumors, 3 µg of total RNA was used for fragmentation without bacterial spike-in RNA. This fragmented RNA was either used for RNA-seq or m^6^A RIP-seq. m^6^A RIP was performed as previously described (5,19,25) with m^6^A antibody. Input and m^6^A RIP RNA were used to generate sequencing libraries using SMARTer Stranded Total RNA-Seq kit V2, Pico Input Mammalian (Takara Bio). All the libraries were single-end sequenced (1×88 bp) on the Illumina NextSeq 2000 platform at the BEA Core Facility (Stockholm, Sweden). Details of the RNA-seq and m^6^A RIP-seq samples used in the study are listed in Supplementary Table S2.

### RT-qPCR and m^6^A RIP-qPCR

For m^6^A RIP-qPCR, 3 µg of cellular RNA was utilized. RIP assays were conducted using 1 µg of m^6^A antibody or IgG antibody as described above. Both input and m^6^A RIP-RNA were reverse-transcribed to cDNA employing the High-Capacity RNA-to-cDNA kit (ThermoFisher) with random primers. Subsequent qPCR was performed on a Quant Studio 3 instrument (ThermoFisher), utilizing gene-specific PCR primers mixed with Power SYBR Green Master Mix (ThermoFisher) and diluted cDNA as a template. The resulting data were expressed as percentage input values. For RT-qPCR, RNA was directly converted into cDNA and subjected to qPCR. The expression values for each gene were normalized to *GAPDH* using the delta-delta Ct method. Primers are listed in Supplementary Table S1.

### RNA stability assay

Transcriptional inhibitor actinomycin D (10 µg/ml) was used to inhibit RNA synthesis. After treatment with actinomycin D, cells were harvested at 3, and 6 h time points, and the RNA was then extracted. The mRNA levels of *HOXC8* and *HOXC9* were detected through RT-qPCR.

### ChIP-seq

ChIP was performed as described before (26). In brief, cells were fixed with formaldehyde (1% final concentration) for 15 min at RT and quenched using Glycine. After fixation, the cells were subjected to cold PBS wash, followed by lysis, and chromatin shearing using a Bioruptor (Diagenode). The chromatin was sheared until the fragments reached an average size range of 200-500 bp. Subsequently, immunoprecipitation of the solubilized chromatin was conducted using 3 µg of METTL3, MYCN, or H3K27ac antibodies overnight at 4°C. The immunoprecipitated complex was then captured using a combination of Protein A and G Dynabeads (Invitrogen), washed and RNase A treated. The samples were next incubated at 68°C for at least 4 h to reverse the cross-links and further treated with proteinase K for 2 h at 37°C. Finally, the ChIP DNA was eluted using the ChIP DNA Clean & Concentrator™ kit (Zymo research, D5205). ChIP-seq libraries were prepared using the NEBNext Ultra II DNA Library Prep Kit (NEB, E7645L) and single-end (1 × 75 bp) sequenced on Illumina NextSeq platform at BEA core facility, Stockholm. Details of the ChIP-seq samples used in the study are listed in Supplementary Table S2.

### Analysis of RNA-seq data

Paired-end RNA-seq reads obtained from BGI DNBseq were analyzed using FastQC v0.11.9 for quality control (https://www.bioinformatics.babraham.ac.uk/projects/fastqc/) using default quality filtering parameter (-q 20). Adapters were removed using Trim Galore v0.6.6 with a minimal length threshold of 20bp. Trimmed reads were mapped to the GRCh38 human reference genome obtained from GENCODE (Release 36 GRCh38.p13) using HISAT2 v2.2.1 (27) with parameters (--sensitive --no-discordant --no-mixed -I 1 -X 1000). After mapping, duplicate alignments were labeled using *markDuplicates* from Picard v2.23.4. Marked alignment files were further processed using Sambamba v0.7.1 (28) keeping uniquely mapping reads separated by strand after duplicate removal. Aligned reads were quantified using Salmon v1.4.0 (29) with GRCh38 Gencode v36 annotation, following differential expression analysis with DESeq2 (30) using two replicates per condition. Genes were considered differentially expressed if their |log2 fold change| > 1 and adjusted p-value < 0.01. Normalized TPM counts were calculated to account for differences in gene length and library sizes for downstream analysis and visualization. Functional enrichment analysis for differentially expressed genes was performed using clusterProfiler (31) and enrichR (32) packages. Data visualization was carried out using custom scripts with the ggplot2 R package (33).

### Analysis of m^6^A RIP data

Single-end m^6^A RIP sequencing data obtained from SMARTer-Stranded Total RNA-Seq Kit v2 was processed using FastQC v0.11.9 and Trim Galore v0.6.6 for quality control. Trimmed reads were mapped to the GRCh38 reference genome using HISAT2 v2.2.1 preserving strand information. Uniquely mapping reads were filtered after duplicate removal and filtering using *markDuplicates* from Picard v2.23.4 and Sambamba v0.7.1, respectively. To control the systematic variation across m^6^A RIP experiments, the amount of spiked-in bacterial RNA was estimated by counting the total number of reads uniquely mapped to the E. coli K-12 reference genome using Sambamba v0.7.1 (28). E. coli spike-in Bacterial counts were further used to calculate scaling factors for each batch of m^6^A RIP-seq samples. Computed scaling factors were then used to normalize the processed alignment files using DownsampleSam tool from Picard v2.23.4. m^6^A modifications were identified following peak calling with MACS2 v2.2.26 (34) on m^6^A RIP and input processed alignments with parameters “--nomodel –bdg –extsize 75 –keep-auto –call-summits” and effective genome size 3.7×10^8^. Called peaks were annotated according to their nearest genomic feature with *annotatePeaks.pl* from HOMER v4.11 (http://homer.ucsd.edu/homer/). Peaks per gene counts were calculated using custom scripts from the annotated peak files. Motif analysis on m^6^A modified peaks was carried out using *findMotifsGenome.pl* from HOMER.

### Metagene analysis

To analyze the genome-wide distribution of m^6^A, a metagene analysis of m^6^A peak density distribution was performed by overlapping the peak coordinates with the genomic features of 5’UTR, CDS, and 3’UTR plus 1 kb upstream and downstream coordinates obtained from GTF genome annotation files from GENCODE v36, the longest isoform for each gene was considered. Each transcript was scaled to fixed size metagene bins according to their respective reference genome coordinates. m^6^A peak density distribution profiles were generated after mapping the m^6^A peaks to the metagene coordinates using the plyranges R package (35). In order to compare multiple conditions, the relative m^6^A density distributions were calculated using the relative density function from ggmulti package (https://cran.r-project.org/web/packages/ggmulti/index.html). The relative density function calculates the sum of the density estimate area of all conditions, where the total sum is scaled to a maximum of 1 and the area of each condition is proportional to its own count.

### Analysis of ChIP-seq data

Single-end METTL3, MYCN, and H3K27ac ChIP-seq data were processed using FastQC v0.11.9 and Trim Galore v0.6.6 for quality control. Trimmed reads were mapped to the GRCh38 reference genome using HISAT2 v2.2.1. Alignment files were further processed using markDuplicates from Picard v2.23.4 and Sambamba v0.7.1 to retrieve mapped reads after duplicate removal. Genome-wide peaks of METTL3, MYCN and H3K27ac ChIP-seq datasets were called using MACS2 v2.2.26 (34) with genome size parameters “-p 1e-5 –nomodel, keep-dup=auto, gsize=2.7e9” for METTL3 ChIP seq data and “-p 1e-9 –nomodel keep-dup=auto, gsize=2.7e9” for MYCN and H3K27ac ChIP-seq datasets as reported in (36,37), respectively. Genome-wide coverage tracks were further calculated using bamCoverage from deepTools v3.3.2 (38). Identified peaks were further annotated according to their nearest genomic feature using annotatePeaks.pl from HOMER v4.11 (http://homer.ucsd.edu/homer/).

To generate Venn diagrams representing the overlap of ChIP-seq peaks, the DiffBind R package was employed (39). The dba.plotVenn function within the DiffBind object, encompassing narrowPeak files along with the corresponding mask, facilitated the calculation of overlapping peaks displayed in the Venn diagram. For the visualization of binding patterns and comparative analysis of raw signals, the ChIPpeakAnno package in R was utilized (40). Specifically, the featureAlignedHeatmap and featureAlignedDistribution functions were employed. To construct the heatmaps, the peaks obtained from the Venn diagram’s overlapping regions were centered on the genomic coordinates of the MYCN obtained from the EnsDb.Hsapiens.v86 R package (41). The peak widths were adjusted and recentred accordingly. Subsequently, the relevant datasets containing BigWig files were processed to create an RleList, which was then utilized to generate the featureAligned signal. The parameter upper.extreme was set to define the upper limit of the color scale, allowing precise control over the visualization of signal intensities. This signal encapsulated the intensity values corresponding to the processed peaks.

### Animal studies

For the *in vivo* experiments, we established tumor xenografts by injecting inducible control (shCtrl) or shMETTL3 SK-N-BE(2) cells subcutaneously into the right dorsal flank of 5-week-old female nude mice (Crl:NU(NCr)-Foxn1nu, Charles River) at a concentration of 5×10^6^ cells in a 200 µL mixture of Matrigel and PBS (1:3 ratio, n=4 per group). To induce METTL3 KD, doxycycline (2 mg/mL) and sucrose (2%) were added to the drinking water approximately 4-5 days after cell injection. We monitored the mice’s weight weekly and measured tumor volume every 2-3 days using a digital caliper, calculated using the formula Volume (mm^3^) = (w2 x l x π)/6, where ‘w’ represents the width (shortest diameter) and ‘l’ represents the length (longest diameter) of the tumor. Mice were euthanized either when the tumors reached 1000 mm^3^ in volume or if they experienced a weight loss of ≥ 10% of their initial weight. Upon conclusion of the experiment, we collected, weighed, and processed the tumors for subsequent analysis. All experiments were carried out as per the standards approved by the Institutional Ethical Committee of Animal Experimentation, Gothenburg, Sweden (ethical permit no 3722/21).

For the drug combination experiments *in vivo*, NSG mice were obtained from Charles River and housed in groups of 2-5 mice per cage in individually ventilated cages with a 12 h light/dark cycle. All procedures were carried out under UK Home Office license P4DBEFF63 according to the Animals (Scientific Procedures) Act 1986 and were approved by the University of Cambridge Animal Welfare and Ethical Review Board (AWERB). COG-N-415x patient-derived xenograft (PDX) cells were obtained from the Childhood Cancer Repository maintained by the Children’s Oncology Group (COG). Cells were suspended in Matrigel (Corning) diluted 1:2 with PBS and 3 × 105 cells (300 µL) were injected into the left flank of NSG mice at an average of 8 weeks of age. Tumors were measured daily with manual calipers and tumor volumes were estimated using the modified ellipsoid formula: V = ab2/2, where a and b (a>b) are length and width measurements respectively. Once tumors reached approximately 170 mm^3^, mice were randomly allocated into four treatment groups (n = 4-6 per group, with the same number of females and males in each study) and treated for 14 days with the following agents by intraperitoneal injection at 10 µl/g body weight: vehicle (20% hydroxypropyl-beta cyclodextrin) daily, STM2457 (50 mg/kg in a vehicle) daily, doxorubicin (0.2 mg/kg in vehicle) every three days or a combination of STM2457 and doxorubicin at the same doses. Mice were euthanized at the end of treatment or once tumors reached 15 mm in any direction (what came first). The maximal tumor size permitted by our Project Licence (15-20 mm) was not exceeded in any of the studies. METTL3 inhibitor STM2457 used in this experiment was synthesized in-house (supplementary methods).

### Statistical analysis

All data were represented as mean ± SD and comparisons between two groups were performed using unpaired two-tailed Student’s *t*-test, and comparisons among more than two groups were performed using one-way or two-way ANOVA (indicated in figure legends). *p* values less than 0.05 were considered statistically significant. The Interaction Factor package in ImageJ was used to randomize the METTL3 signal distribution to investigate the potential effects of random cluster overlap (42). All graphs were generated using GraphPad Prism software (version 10) or R with ggplot2.

## Results

### METTL3 regulates posterior *HOX* genes expression during differentiation of tNCC

We have established a protocol for *in vitro* differentiation of hESC to tNCC adapting the previously described methodology (15) (**Fig. 1A**). The tNCC can be further differentiated into sympathoadrenergic progenitors (SAP) and then into sympathetic neurons (SN). To confirm differentiation, using immuno-fluorescence (IF), robust expression of HOXC9, a posterior *HOX* gene was seen in tNCC whereas SAP cells expressed *PHOX2B,* and the differentiated SN were positive for peripherin (PRPH) (**Fig. 1B**). At the RNA level the pluripotency markers, *OCT4* and *NANOG* decreased during differentiation whereas *HOXC9*, *SOX10,* and *NGFR* were upregulated at the tNCC stage (Supplementary Fig. S1A). In addition, SAP showed upregulation of *ASCL1* and *ISL1* whereas typical SN markers (*DBH* and *TH*) were upregulated from the SAP stage of differentiation onwards with expression maintained in SN (Supplementary Fig. S1A). Global gene expression changes comparing those of hESC and tNCC by RNA-seq showed that differentially expressed genes (DEGs) are enriched with pathways related to anterior-posterior pattern formation, epithelial to mesenchymal transition (EMT), and neural crest differentiation (Supplementary Fig. S1B). Upregulation of *HOX* genes could be also detected in tNCC using RNA-seq data (Supplementary Fig. S1C). Overall, these data suggest that differentiation of hESC to tNCC, SAP, and SN had been achieved in our model system.

**Figure 1.**
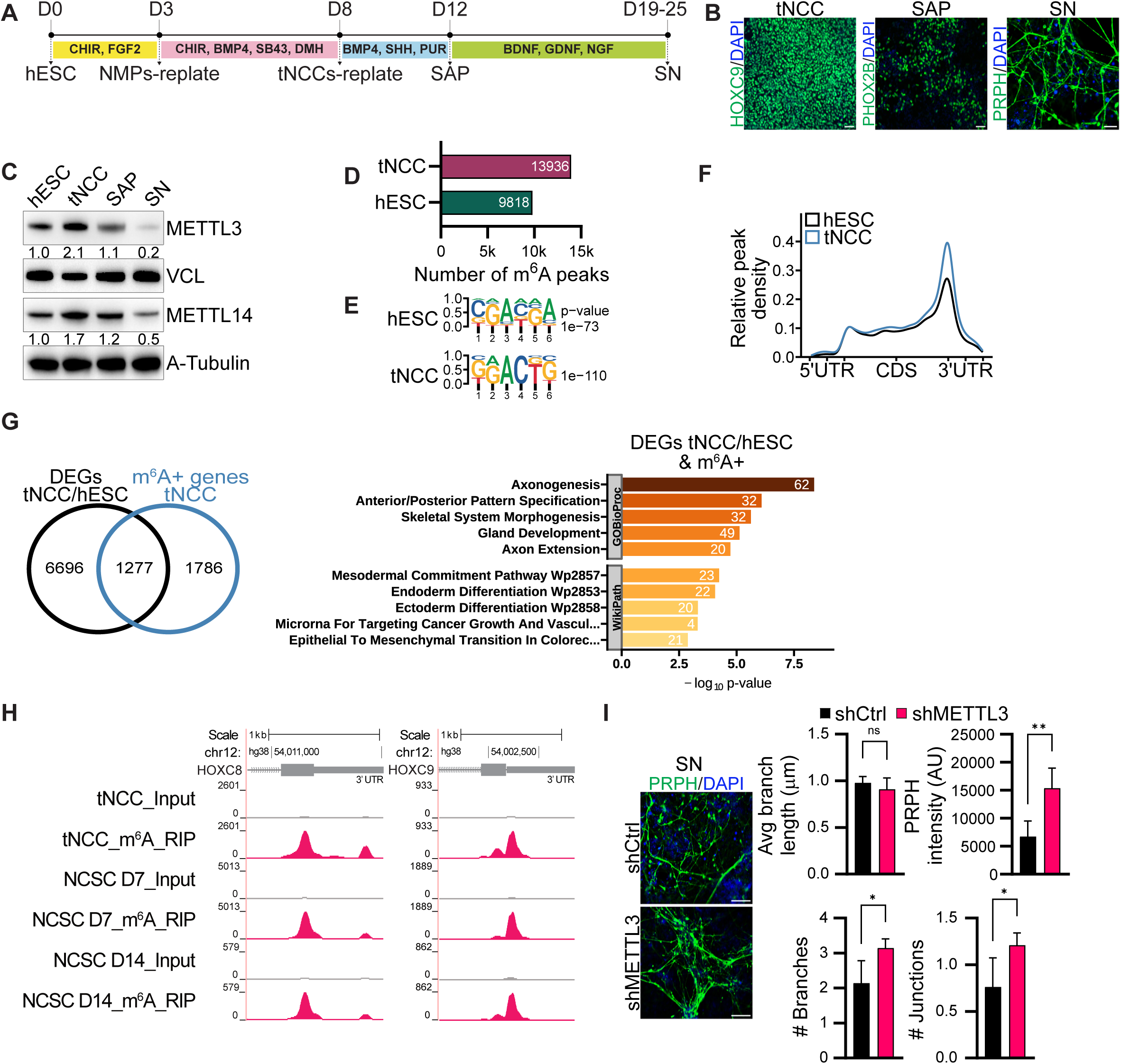
A, Schematic diagram showing key steps involved in the differentiation process of human embryonic stem cells (hESC) into trunk neural crest cells (tNCC), followed by their further differentiation into sympathoadrenal progenitors (SAP) and, ultimately, into sympathetic neurons (SN). B, Representative Immunofluorescence (IF) images illustrating the expression of distinct lineage markers at different stages of differentiation: HOXC9 for trunk neural crest cells (tNCC), PHOX2B for sympathoadrenal progenitors (SAP), and PRPH for sympathetic neurons (SN). The scale bar is indicative of 50 µm. C, Representative immunoblot shows the levels of METTL3 and METTL14 across various stages of differentiation, including hESC, tNCC, SAP, and SN. Vinculin and A-tubulin were loading controls. The values below the blots indicate the fold change in the levels of METTL3 or METTL14. D, The total number of m6A peaks in hESC and tNCC. E, Identified motifs from de novo motif analysis of m6A peaks enriched in hESC and tNCC. F, Metagene analysis showing relative m6A peak density at genes in hESC and tNCC. G, (Left panel) Venn diagram comparison of differentially expressed genes (DEGs) [hESC vs. tNCC] and m6A positive [(m6A +) having at least one m6A peak] genes in tNCC. (Right panel) Top enriched terms associated with DEGs (hESC vs. tNCC) having m6A. H, Genome browser screenshots of HOXC8 and HOXC9 3′UTR, showing the presence of m6A in tNCC, neural crest stem cells (NCSC) at day 7 and at day 14. I, Control (shCtrl), and stable METTL3 KD (shMETTL3) hESC were differentiated to SN followed by IF with PRPH antibody to assess PRPH signal intensity, neurite branch length, as well as the number of branches and junctions. The quantification results are depicted in bar plots. Data are shown as mean ± SD and this analysis was conducted across three independent biological replicates. Statistical significance was determined using an unpaired t-test (* p < 0.05, ** p < 0.01, ns - non-significant p > 0.05). Scale bar represents 100 µm.

Having established cellular differentiation, we sought to examine expression levels of METTL3 and METTL14 proteins which reached a peak at the tNCC stage and then gradually decreased as the cells transitioned through the SAP to SN stage (**Fig. 1C**). However, the regulation of expression of these proteins at the hESC to tNCC stage is likely regulated post-transcriptionally as RNA levels of *METTL3* and *METTL14* were unchanged (Supplementary Fig. S1D). Indeed, METTL3 was more stable at the tNCC stage as shown by a cycloheximide chase experiment conducted with cells at both the hESC and tNCC stages of differentiation (Supplementary Fig. S1E).

Next, we characterized the pattern of m^6^A modifications in hESC and tNCC using m^6^A RIP-seq (19). We observed that consistent with the upregulation of METTL3/14, a higher number of m^6^A peaks was seen in tNCC compared to hESC (**Fig. 1D**), and these were enriched with DRACH like motifs (**Fig. 1E**). tNCC showed a higher number of m^6^A peaks and a higher relative m^6^A peak density in comparison with hESC indicating a potentially important role for m^6^A modification in differentiation of tNCC (**Fig. 1F**).

The top enriched terms of the genes associated with m^6^A peaks from both hESC and tNCC showed pathways related to RNA splicing consistent with the role of m^6^A modification in RNA metabolism (Supplementary Fig. S1F). Furthermore, we observed that the genes that were modified by m^6^A in tNCC and differentially expressed between hESC and tNCC were enriched in pathways related to anterior-posterior pattern formation, axonogenesis, and EMT (**Fig. 1G**). Overall, higher expression of METTL3/14, differential gene expression along with m^6^A RIP-seq data in hESC and tNCC suggest a critical role for m^6^A in tNCC.

To validate the role of m^6^A modification in neural crest cell differentiation we used an alternative previously described protocol to generate multipotent neural crest stem cells (NCSC) from hESC (20). The NCSC identity of these cells was validated by robust expression of neural crest lineage markers (Supplementary Fig. S1G). During the NCSC differentiation protocol, the cells were harvested at different stages of differentiation (days 3, 7, and 14), METTL3 and METTL14 expression were upregulated during NCSC differentiation compared to hESC (Supplementary Fig. S1H). Consistent with the increase in the METTL3 level at day 3 NCSC progenitors, the stability of the METTL3 protein was higher compared to hESC (Supplementary Fig. S1E). We also performed m^6^A RIP-seq of day 7 NCSC progenitors and NCSC at day 14 and detected 10,713 and 7,250 m^6^A peaks respectively, suggesting a critical role for m^6^A during NCSC differentiation (Supplementary Fig. S1I).

To further characterize the role of METTL3 mediated m^6^A modification in tNCC we generated METTL3 knockdown (KD) hESC which had been differentiated to tNCC or NCSC (Supplementary Fig. S1J). Gene expression analysis of the METTL3 KD tNCC and day 7 NCSC showed robust upregulation of *HOX* genes (Supplementary Fig. S1K). Furthermore, the genes that were deregulated following METTL3 KD and had m^6^A peaks in tNCC were associated with pathways related to anterior-posterior pattern specification (Supplementary Fig. S1L). In particular, the posterior *HOX* genes *HOXC8* and *HOXC9* were enriched with m^6^A in tNCC, day 7 NCSC progenitors, and day 14 NCSC (**Fig. 1H**). RNA-seq data showed METTL3 KD resulted in upregulation of *HOXC8* and *HOXC9* in day 7 NCSC progenitors and *HOXC8* in tNCC (Supplementary Fig. S1K). In addition, METTL3 KD resulted in upregulation of both, *HOXC8* and *HOXC9* as detected by RT-qPCR at SAP suggesting m^6^A dependent regulation of these genes (Supplementary Fig. S1M). METTL3 KD SAP showed deregulation in the expression of several SAP markers such as *PHOX2B*, *ASCL1*, *ISL1*, and *GATA2* suggesting METTL3 KD created a change in the differentiation potential of these cells (Supplementary Fig. S1N). On further differentiation to the SN stage, METTL3 KD promoted higher differentiation to SN as visualized increased PRPH intensity, number of neuronal branching, and junctions, despite no changes observed in neurite branch length (**Fig. 1I**).

In addition, conditional KD of METTL3 (Dox induced KD from day 5 onwards) specifically at the tNCC also led to enhanced differentiation at the SN stage as visualized by increased PRPH intensity (Supplementary Fig. S1O and S1P). These data suggest that the differentiation phenotype we observed is not due to METTL3 KD at the hESC stage but rather a reduced level of METTL3 during the differentiation of tNCC. Conditional METTL3 KD at the tNCC stage also led to the upregulation of HOXC8 and HOXC9 expression in the differentiated SAP (Supplementary Fig. S1Q). Our data suggest METTL3 has a critical role in regulating the timely transition of tNCC to SAP through regulating the expression of posterior *HOX* genes such as *HOXC8* and *HOXC9* via m^6^A modification.

### METTL3 mediated m^6^A modification controls HOXC8 and HOXC9 expression in MYCN-amplified NB tumors and in NB cell lines

Using RNA-seq data from 498 NB tumor samples we observed low expression of *HOXC8* and *HOXC9* correlates with poor survival and this is consistent with an earlier report (**Fig. 2A**) (43). We also observed that *HOXC8* and *HOXC9* expression was downregulated in MYCN-amplified tumors **(Fig. 2B)**. To explore this further, we performed m^6^A RIP-seq of RNA derived from MYCN-amplified NB tumors and observed m^6^A modification in *HOXC8* and *HOXC9* transcripts (**Fig. 2C**). The m^6^A peaks in the NB tumors were enriched at the stop codon and 3′UTR regions and with DRACH like motifs as reported earlier (44) (Supplementary Fig. S2A). The common m^6^A enriched genes detected in MYCN-amplified tumors were related to pathways such as axonogenesis and dendrite development/morphogenesis (**Fig. 2D)**. These data were further enforced by analyzing m^6^A RIP-seq of MYCN-amplified SK-N-BE(2) cells in which *HOXC8*/*HOXC9* were also modified by m^6^A (**Fig. 2E**). Next, we examined whether the depletion of METTL3 in SK-N-BE(2) cells leads to the upregulation of *HOXC8* and *HOXC9* similarly. Stable shRNA mediated depletion of METTL3 was not possible with repeated attempts as METTL3 depleted cells did not survive, suggesting METTL3 was essential for the survival of SK-N-BE(2) cells. Hence, a doxycycline (Dox) inducible shRNA system was employed to deplete METTL3, and cells were analyzed by RNA-seq. These data showed posterior *HOXC* locus genes such as *HOXC8*, *HOXC9,* and *HOXC10* were upregulated and were m^6^A modified (**Fig. 2E** and **F**). In particular, we observed the increased stability of *HOXC8* and *HOXC9* mRNA following METTL3 KD in SK-N-BE(2) cells (**Fig. 2G).** Furthermore, Dox induced METTL3 KD in SK-N-BE(2) and IMR-32 (both MYCN-amplified NB cell lines) led to reduced proliferation in both cases (Supplementary Fig. 2B). To further verify the effects of METTL3 KD in combination with MYCN overexpression we have used SHEP cells (low MYCN expressing NB cells), using Dox inducible system. We observed that METTL3 KD in MYCN overexpressed SHEP cells decreased cell viability (Supplementary Fig. 2C). Moreover, injection of Dox inducible METTL3 KD SK-N-BE(2) cells into immunocompromised nude mice administered with doxycycline, led to reduced xenografted tumor growth (**Fig. 2H**). We validated METTL3 KD in xenografted SK-N-BE-2 tumors with consequent upregulation in HOXC9 expression (**Fig. 2H)**. Interestingly, METTL3 KD resulted in downregulation of METTL14 expression as well (**Fig. 2H)**. We verified a decrease in METTL14 expression by siRNA mediated KD of METTL3 in SK-N-BE(2) cells (Supplementary Fig. S2D). This observation is consistent with a recent report that suggested METTL3 protects METTL14 by preventing its ubiquitination and degradation (11,45). METTL3 KD also reduced colony formation in SK-N-BE(2), in line with our *in vivo* findings (Supplementary Fig. S2E; **Fig. 2H**). The genes that were deregulated after METTL3 KD and m^6^A positive in SK-N-BE(2) cells were enriched in pathways related to axonogenesis suggesting m^6^A dependent role of METTL3 in the differentiation of NB cells (Supplementary Fig. S2F). To explore this further, we have performed RA mediated differentiation of control and Dox inducible METTL3 KD SK-N-BE(2) cells and we observed METTL3 KD could promote differentiation of SK-N-BE(2) cells (**Fig. 2I**). Stable overexpression of both *HOXC8* and *HOXC9* induced spontaneous differentiation of SK-N-BE(2) cells without RA addition (Supplementary Fig. S2G) suggesting upregulation of posterior *HOX* genes in the METTL3 depleted cells could be a major driver for differentiation (**Fig. 2I)** (43,46). We found around 20% of *HOXC9* target genes identified by ChIP-seq (46) overlapped with METTL3 deregulated genes. These overlapped genes with a role in neuronal differentiation such as PRPH further suggesting *HOXC9* upregulation in METTL3 KD cells contributes to the differentiation phenotype we observed (Supplementary Fig. S2H).

**Figure 2.**
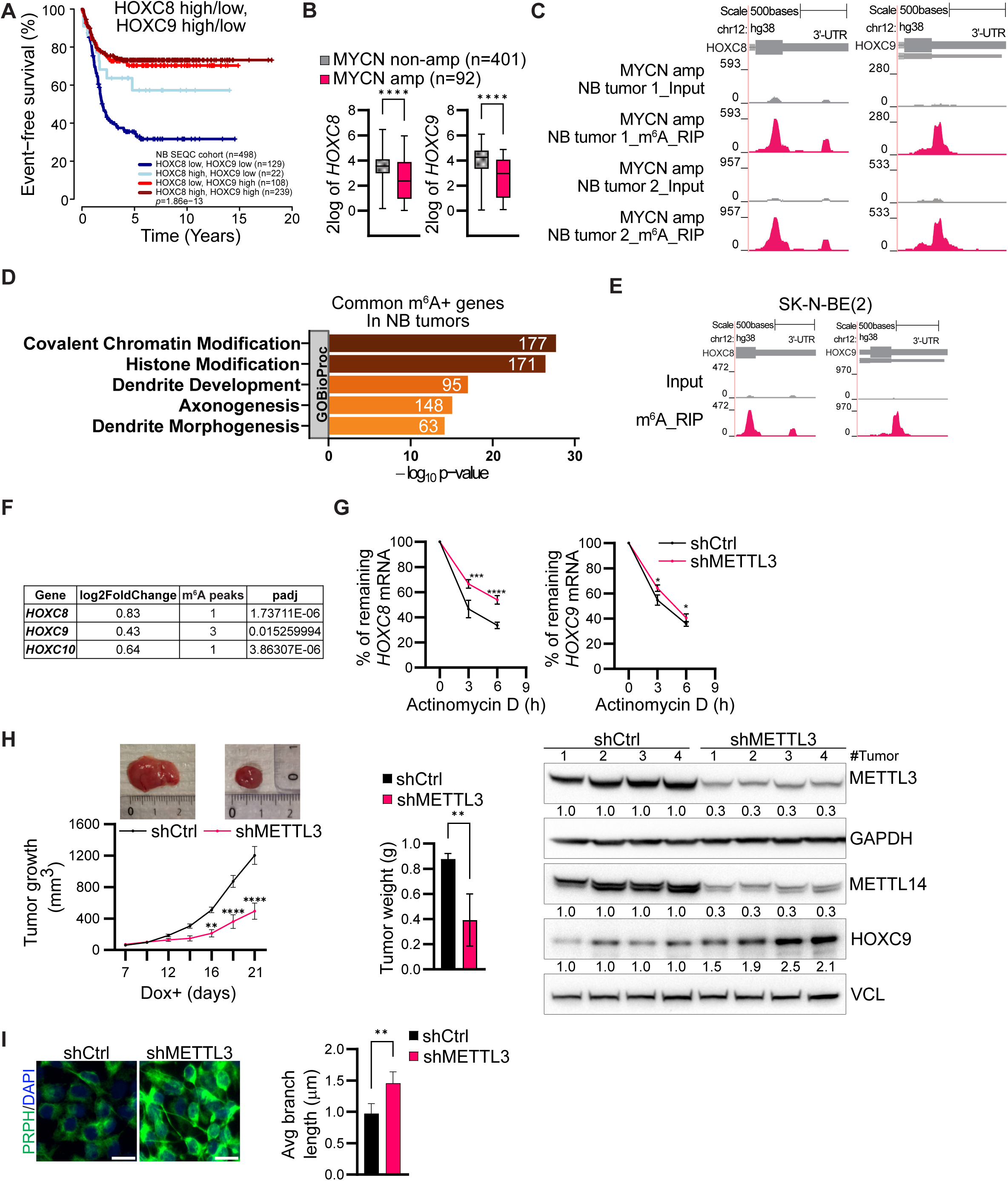
A, Kaplan-Meier plot illustrates event-free survival in neuroblastoma (NB) patients (n=498, SEQC cohort) with either low or high expression of HOXC8 and HOXC9. Statistical analysis of survival was performed with a log-rank test. B, Box and whisker plots show HOXC8 and HOXC9 expression in NB patients (SEQC cohort) classified based on MYCN amplification status. The centrelines of the box are medians, and the upper and lower lines indicate the 25th and 75th percentiles. Statistical analysis was performed using two-sided Mann–Whitney tests (**** p < 0.0001). C, Browser screenshot showing the presence of m6A at 3′UTR of HOXC8 and HOXC9 genes in MYCN-amplified NB tumors. D, Top enriched terms associated with m6A+ genes in both NB tumors. E, Genome browser screenshot showing the presence of m6A enrichment at 3′UTR of HOXC8 and HOXC9 genes in MYCN-amplified NB cell line SK-N-BE(2). F, Differentially expressed posterior HOXC genes between control and METTL3 KD SK-N-BE(2) cells and the number of m6A peaks identified using MACS peak caller in these genes are indicated. G, Stability of HOXC8 and HOXC9 transcripts detected by RT-qPCR post Actinomycin D (10 µg/ml) mediated transcription blocking for the time points indicated in METTL3 KD SK-N-BE(2). Line plots presenting the quantification of remaining levels of HOXC8 and HOXC9 transcript at the indicated time points. Two-way ANOVA with Šídák’s multiple comparisons test was employed (* p < 0.05, *** p < 0.001, **** p < 0.0001). H, (Left panel) Line plots showing tumor volume in Control and METTL3 KD SK-N-BE(2) mouse xenograft with representative tumors from each group (n=4). (Middle panel) Bar plot shows tumor weight in Control and METTL3 KD xenograft tumors. (Right panel) Immunoblot showing expression of METTL3, METTL14, and HOXC9 in Control and METTL3 KD xenografted tumors. GAPDH and vinculin were used as loading controls. The values below indicate the fold change in levels of METTL3, METTL14, and HOXC9. Two-way ANOVA with Šídák’s multiple comparisons test was employed to compare tumor volumes and unpaired t-test for tumor weights (** p < 0.01, **** p < 0.0001). I, Representative IF showing PRPH (green) staining in Control and METTL3 KD SK-N-BE(2) cells following 3 days RA mediated differentiation. Bar plot shows the quantification of the neurite branch length. Scale bar represents 5 µm. Experiments were performed in three independent biological replicates. Unpaired t-test was used, (** p < 0.01).

Network analysis predicted HOXC8 and HOXC9 could have functional interaction and our proximity ligation assay (PLA) data showed *HOXC8* and *HOXC9* interaction in MYCN-amplified NB cells (Supplementary Fig. S2I). The interaction between HOXC8 and HOXC9 explains the stronger RA mediated differentiation phenotype we observed after transient overexpression of both the *HOXC8* and *HOXC9* genes together compared to a single *HOX* gene in SK-N-BE(2) cells (Supplementary Fig. S2J).

### MYCN overexpression creates an undifferentiated state in tNCC

Interestingly, MYCN expression is high in hESC and tNCC but is then downregulated as differentiation progresses to SAP, and it becomes almost undetectable in SN cells (**Fig. 3A**). Given that MYCN is amplified in NB, our tNCC to SN differentiation model allows us to explore how MYCN deregulation may contribute to the improper differentiation characteristic of NB. To enforce MYCN expression throughout differentiation, we created a Dox inducible expression system by introducing MYCN into hESC using the inducible PiggyBac system (21). To induce the MYCN overexpression Dox was added from day 5 of differentiation and continued until the end of the experiment (**Fig. 3B**). We validated the overexpression of MYCN following induction with Dox at the tNCC and SAP stages of differentiation by immunoblot (**Fig. 3B** and **C**). To determine whether MYCN overexpression affects differentiation towards the SN stage, we harvested cells at day 22 when the SN phenotype is normally observed and noted that the cells forcibly expressing MYCN lacked expression of PRPH and TUBB3 as detected by IF (**Fig. 3D** and **E**). We wanted to check if HOXC8 and HOXC9 expression was altered following MYCN overexpression, and expression was altered following MYCN overexpression, and we observed HOXC8 and HOXC9 were downregulated in MYCN overexpressing SAP cells (**Fig. 3F** and **G**; Supplementary Fig. S3A). The stability of *HOXC8* and *HOXC9* transcripts was also reduced in MYCN overexpressed SAP stage cells (Supplementary Fig. S3B). To further investigate transcriptional changes on MYCN overexpression at the SN stage, we conducted RNA-seq. Consistent with the undifferentiated phenotype (**Fig. 3D** and **E**) we observed that downregulated genes were enriched in pathways related to axonogenesis, axon guidance, and neuronal projection guidance (Supplementary Fig. S3C). In contrast, upregulated genes following MYCN overexpression in SN were related to the RNA metabolic process, which is consistent with a role for MYCN in global transcriptional upregulation as previously observed (Supplementary Fig. S3C)(47).

**Figure 3.**
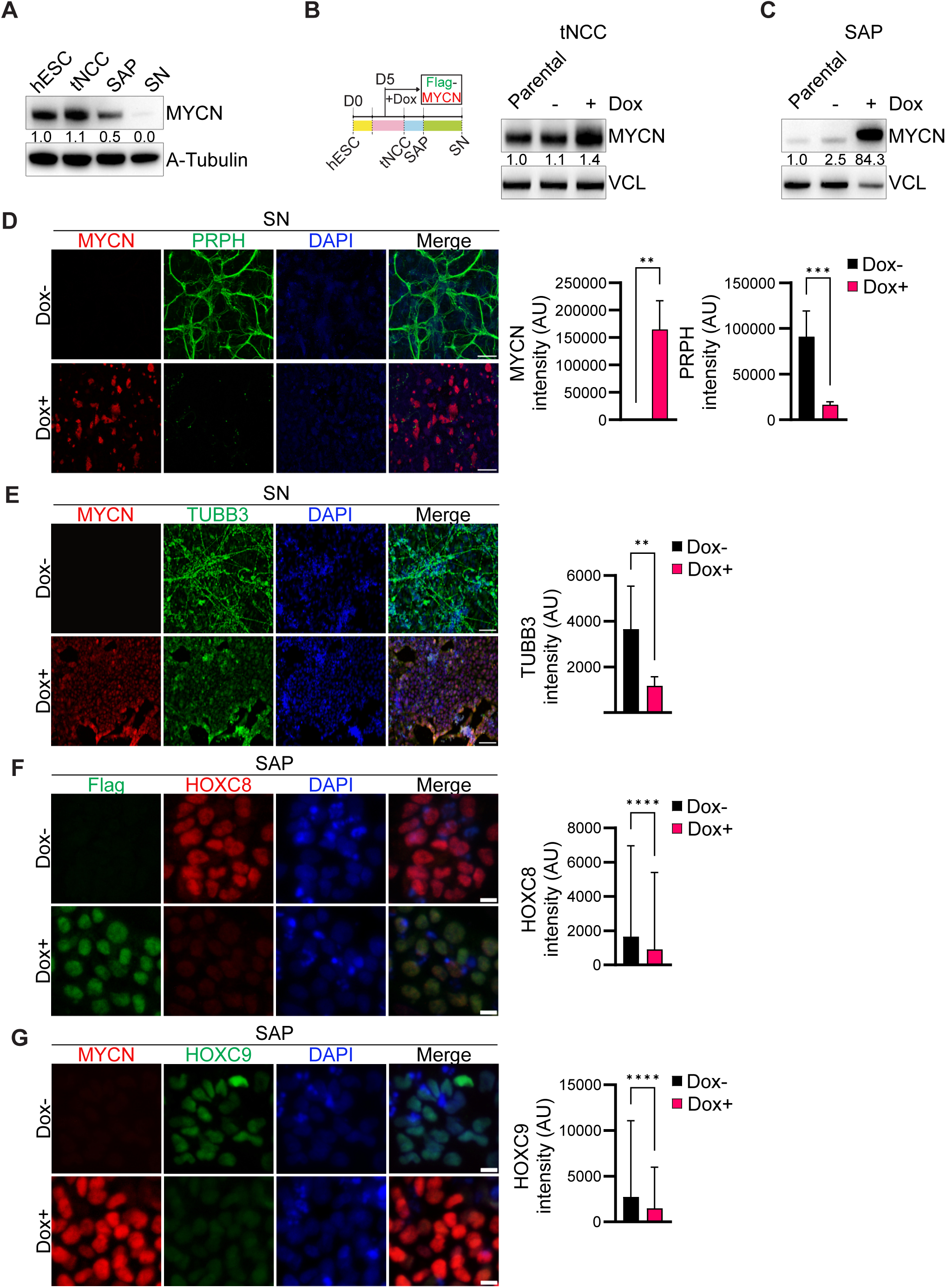
A, Representative immunoblot showing expression of MYCN in hESC, tNCC, SAP, and SN stages. A-tubulin was used as a loading control. The values below indicate the fold change in levels of MYCN. The experiments were repeated three times. B, (Left panel) Schematic diagram showing time of FLAG-tagged MYCN (Flag-MYCN) induction in Dox dependent manner from day 5 during tNCC differentiation. (Right panel) Immunoblot showing MYCN overexpression in tNCC. Vinculin was used as a loading control. The values below indicate the fold change in levels of MYCN. C, Immunoblot showing the level of MYCN expression in SAP following Dox induced Flag-MYCN from day 5 of differentiation. Vinculin was used as a loading control. The values below indicate the fold change in levels of MYCN. D, Control (Dox-) and Flag-MYCN overexpressed (Dox+, from day 5 onwards) tNCC were differentiated to SN and IF was performed with PRPH (green) and MYCN (red) antibodies. Bar plots show MYCN and PRPH signal intensity. Data are presented as mean ± SD from three independent experiments. Statistical analysis was performed using an unpaired t-test (** p < 0.01, *** p < 0.001). Scale bar represents 500 μm. E, Control (Dox-) and Flag-MYCN overexpressed (Dox+, from day 5 onwards) tNCC were differentiated to SN and IF was performed with TUBB3 (green) and MYCN (red) antibodies. Bar plot shows TUBB3 intensity. Data are presented as mean ± SD from three independent experiments. Statistical analysis was performed using an unpaired t-test (** p < 0.01). Scale bar represents 100 μm. F, Representative IF showing expression of HOXC8 (red) in control (Dox-) and Flag-MYCN overexpressed (Dox+, from day 5 onwards) SAP. MYCN (green) was visualized with an anti-FLAG antibody. Bar plot shows HOXC8 signal intensity. Signal intensity measurements were taken from over 2000 cells. Data are presented as mean ± SD from three independent experiments. Statistical analysis was performed using an unpaired t-test (**** p < 0.0001). Scale bar represents 10 μm. G, Representative IF showing expression of HOXC9 (green) in control (Dox-) and Flag-MYCN overexpressed (Dox+, from day 5 onwards) SAP. MYCN (red) was visualized with MYCN antibody. Bar plot shows HOXC9 intensity. Signal intensity measurements were taken from over 2000 cells. Data are presented as mean ± SD from three independent experiments. Statistical analysis was performed using an unpaired t-test (**** p < 0.0001). Scale bar represents 10 μm.

### MYCN cooperates with METTL3 to regulate m^6^A levels of HOXC8 and HOXC9

Our data shows that the alteration in MYCN and METTL3 levels could regulate HOXC8 and HOXC9 expression in the tNCC differentiation system (**Fig. 3D-E** and **1I)**. This suggests that MYCN might cooperate with METTL3 to regulate m6A levels during tNCC differentiation and thereby gene expression. To explore this further, we performed Co-immunoprecipitation (Co-IP) with MYCN antibody in SHEP cells following Dox induced MYCN overexpression and found METTL3 to be interacting with MYCN. This was also true vice-versa, i.e. when METTL3 was immunoprecipitated and MYCN was immunoblotted (Fig. 4A). We further performed PLA and observed an interaction between MYCN and METTL3 in tNCC (Fig. 4A). In addition, in SHEP cells following Dox induced MYCN overexpression (Supplementary Fig. S2C) an interaction between MYCN and METTL3 was also observed by PLA (Fig. 4A). Furthermore, IF for MYCN and METTL3 in tNCC showed co-localization of these proteins in the cells (**Fig. 4B**).

**Figure 4.**
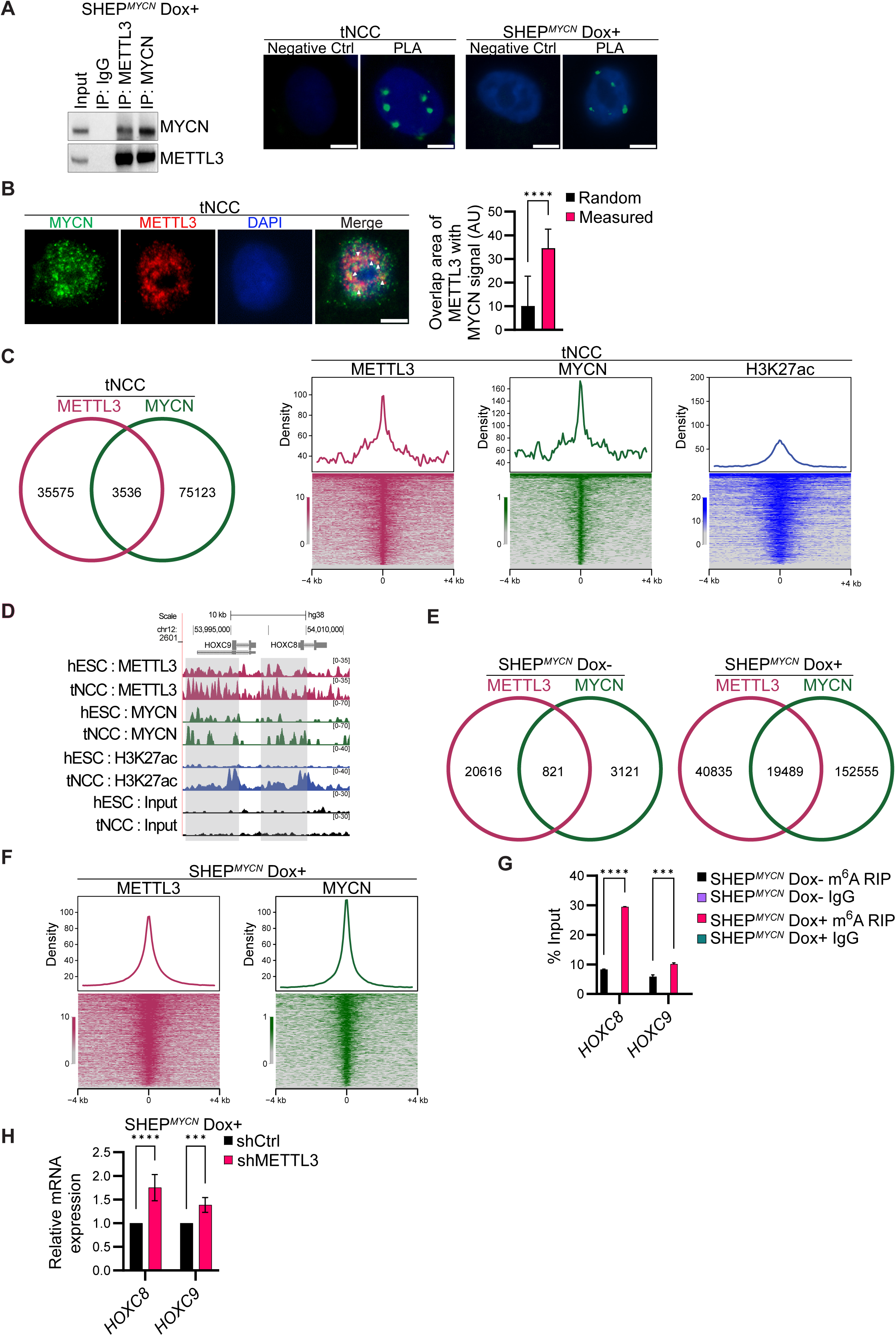
A, (Left panel) Co-IP of METTL3 or MYCN from lysates of SHEPMYCN after Dox induction for 24 h, blotted with MYCN or METTL3 antibodies. IgG served as negative control. (Right panel) Proximity ligation assay (PLA) in tNCC and 24 h Dox induced SHEPMYCN cells depicting METTL3 and MYCN PLA signal (green) in the nucleus (marked by DAPI). The negative control shows PLA with only the METTL3 antibody. Scale bar is 50 μm. B, (Left panel) MYCN (green) and METTL3 (red) IF was performed in tNCC. White arrows indicate the colocalization of METTL3 and MYCN. (Right panel) Bar plot shows the overlap area of METTL3 with MYCN. The area of the overlapping METTL3 and MYCN signal was quantified using the Interaction Factor package in ImageJ. To assess the significance of the observed overlapped area, the METTL3 signal was randomized for each image. The means of 20 randomizations were then plotted alongside the experimentally observed values. This analysis allowed us to evaluate the statistical significance of the observed overlap between METTL3 and MYCN signals. At least 50 cells were counted from two independent biological replicates. Unpaired t-test was used, **** p < 0.0001. C, (Left panel) Venn diagram comparison of METTL3 and MYCN binding sites determined from ChIP-seq experiments performed in tNCC. (Right panel) Distribution and heatmaps of normalized ChIP–seq reads for METTL3, MYCN, and H3K27ac over the MYCN and METTL3 overlapping peak coordinates. The data is centered on MYCN peaks. D, Genome browser screenshot showing METTL3, MYCN, and H3K27ac ChIP-seq signals in hESC and tNCC over the HOXC8 and HOXC9 gene locus. E, Venn diagram comparison of METTL3 and MYCN binding sites determined from ChIP-seq experiments performed in SHEPMYCN cells before and after Dox induction for 24 h. F, Distribution, and heatmaps of normalized ChIP–seq reads for METTL3 and MYCN overlapping peaks centered on MYCN peaks in SHEPMYCN cells after Dox induction. G, m6A RIP-qPCR data showing enrichment of both HOXC8 and HOXC9 in SHEPMYCN cells before and after Dox induction for 24 h. Data are represented as a percentage of input. IgG was used as a negative control (n=3). Two-tailed paired t-test was used, *** p < 0.001, **** p < 0.0001. H, RT-qPCR data showing the expression of HOXC8 and HOXC9 in SHEPMYCN cells with either Control (shCtrl) or stable METTL3 KD (shMETTL3) after Dox induction for 6 days. GAPDH was used to normalize the qPCR data. Data are shown as mean ± SD of three replicates. Two-tailed paired t-test was used, *** p < 0.001, **** p < 0.0001.

The mechanisms by which METTL3 is recruited to chromatin are largely unknown. Our data whereby MYCN co-localizes with METTL3 in the nucleus (**Fig. 4A** and **B**) suggests that MYCN might guide METTL3 to distinct genomic regions. To explore this further, ChIP-seq of both MYCN and METTL3 was conducted in hESC and tNCC. We observed an overlap between MYCN and METTL3 chromatin-bound regions genome-wide in both hESC and tNCC (Supplementary Fig. S4A; **Fig. 4C**). The MYCN and METTL3 co-bound regions were enriched with active chromatin modification H3K27ac in tNCC and were located around gene promoters (**Fig. 4C**; Supplementary Fig. S4B and S4C).

Approximately 20% of MYCN and METTL3 promoter bound genes in tNCC contained at least one m^6^A peak (Supplementary Fig. S4D). Interestingly, the MYCN and METTL3 promoter bound and m^6^A modified genes which were deregulated upon METTL3 KD in tNCC were related to axonogenesis (Supplementary Fig. S4D). Furthermore, higher enrichment of METTL3 and MYCN was seen over the *HOXC8* and *HOXC9* gene loci in tNCC compared to hESC, and these genes were also enriched with the active histone modification H3K27ac in tNCC but not hESC, consistently with their expression at this stage of differentiation (**Fig. 4D**). To determine whether MYCN expression can influence METTL3 binding to genomic regions, we utilized the Dox inducible SHEP*^MYCN^* system (Supplementary Fig. 2C). Mapping of METTL3 and MYCN binding before and after MYCN overexpression in SHEP cells showed an expected increase in MYCN binding genome-wide on Dox induction (**Fig. 4E**). In addition, a 3-fold increase in the number of METTL3 peaks was also observed in comparison with MYCN non-induced (Dox-) SHEP cells (**Fig. 4E**). MYCN and METTL3 binding sites frequently overlapped in Dox induced SHEP*^MYCN^* cells and again most of the overlapping peaks were associated with gene promoters (**Fig. 4E** and **F**; Supplementary Fig. S4E). Furthermore, MYCN overexpression resulted in increased METTL3 recruitment to HOXC8 and HOXC9 genes (Supplementary Fig. S4F), and this correlated with a higher level of m^6^A enrichment at these genes as detected by m^6^A RIP-qPCR (**Fig. 4G**). Consistent with this, combined MYCN overexpression and METTL3 KD resulted in increased expression of *HOXC8* and *HOXC9* compared to MYCN overexpression alone (**Fig 4H**).

METTL3 has recently been reported to regulate global transcription by upregulating MYCN expression in paused mouse ES cells (48). However, we did not observe such an effect of METTL3 KD on MYCN expression levels in tNCC or NB cells (Supplementary Fig. S4G). Our data is consistent with other observation (49) in NB cells where METTL3 KD did not impact MYCN expression levels (Supplementary Fig. S4G).

### METTL3 inhibition restores differentiation and sensitizes NB cells to chemotherapeutic drug

Our data show that MYCN overexpression downregulates and METTL3 KD upregulates expression of the posterior *HOX* genes *HOXC8* and *HOXC9* suggesting potential antagonistic regulation (Supplementary Fig. S1M and S3A; **Fig. 3F** and **G**). On the other hand, *HOXC8* and *HOXC9* over-expression promoted differentiation of the MYCN-amplified SK-N-BE(2) cells (Supplementary Fig. S2G and S2J). Hence, we proposed that inhibition of METTL3 with a small molecule inhibitor could promote differentiation of MYCN overexpressing tNCC. We treated MYCN overexpressing tNCC with STM2457, a small molecule inhibitor of METTL3, recently developed as a therapeutic strategy for acute myeloid leukemia (AML) (18). HOXC8 and HOXC9 expression was restored in STM2457 treated MYCN overexpressing SAP stage cells (**Fig. 5A** and **B).** Indeed, long term treatment of MYCN overexpressing tNCC with STM2457 rescued the differentiation phenotype of these cells as observed by PRPH IF (**Fig. 5C**). To verify the specificity of STM2457 mediated METTL3 inhibition induced rescue of the differentiation phenotype in MYCN overexpressing tNCC, we simultaneously knocked down METTL3 (from day 5 onwards) in the MYCN overexpressing tNCC (Supplementary Fig. S5A). As expected, METTL3 KD rescued the differentiation of MYCN overexpressing cells as evidenced by PRPH and TUBB3 expression detected by IF (Supplementary Fig. S5A). Hence, our data suggest that MYCN co-operates with METTL3 to create an undifferentiated state that can be reversed by METTL3 inhibition or KD.

**Figure 5.**
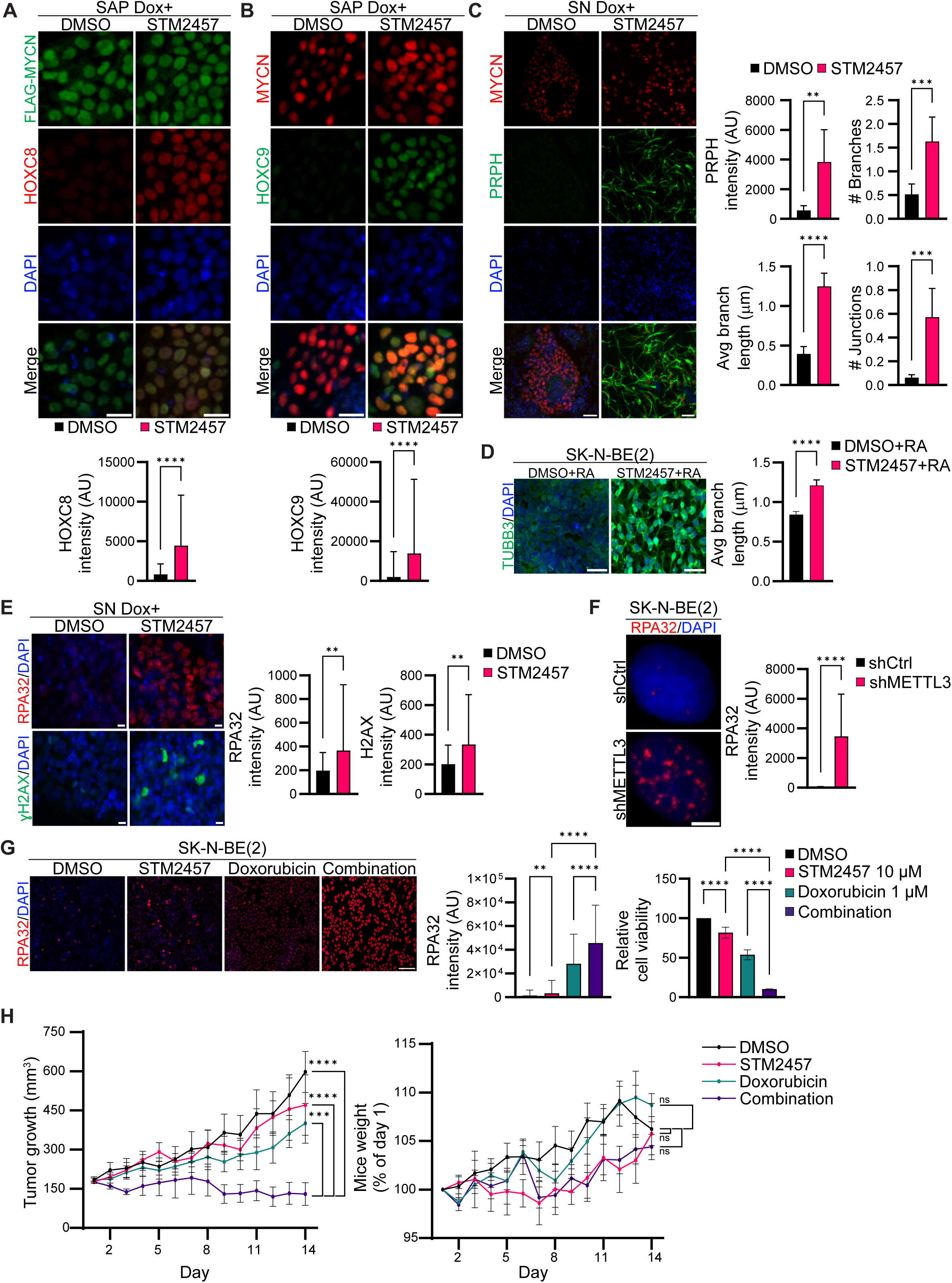
A, (Top panel) HOXC8 (red) and Flag (green) IF was performed in Flag-MYCN overexpressed (Dox+, from day 5 onwards) SAP after DMSO or STM2457 (10 μM) treatment. STM2457 or DMSO was added on day 9 of differentiation. (Bottom panel) Bar plot shows HOXC8 signal intensity. Signal intensity measurements were taken from over 1000 cells and data are presented as mean ± SD from three independent experiments. Statistical analysis was performed using an unpaired t-test (**** p < 0.0001). Scale bar represents 10 μm. B, (Top panel) HOXC9 (green) and MYCN (red) IF was performed in Flag-MYCN overexpressed (Dox+, from day 5 onwards) SAP after DMSO or STM2457 (10 μM) treatment. STM2457 or DMSO was added on day 9 of differentiation. (Bottom panel) Bar plot shows HOXC9 intensity. Signal intensity measurements were taken from over 1000 cells and data are presented as mean ± SD from three independent experiments. Statistical analysis was performed using an unpaired t-test (**** p< 0.0001). Scale bar represents 10 μm. C, (Left panel) PRPH (green) and MYCN (red) IF was performed in Flag-MYCN overexpressed (Dox+, from day 5 onwards) SN stage cells after DMSO or STM2457 (10 μM) treatment. STM2457 or DMSO was added from day 9 of differentiation. (Right panel) Bar graph shows either PRPH signal intensity, quantification of neurite branch length, number of branches, and junctions. Data are presented as mean ± SD from three independent experiments. Statistical analysis was performed using an unpaired t-test (** p < 0.01, *** p < 0.001, **** p < 0.0001). Scale bar is 50 μm. D, Representative IF images of TUBB3 (green) in SK-N-BE(2) cells that were pre-treated with either DMSO or STM2457 (10 μM) for 24 h, followed by RA treatment for another 3 days. (Right panel) Bar plot shows the quantification of neurite branch length. Data are presented as mean ± SD from three independent experiments. Statistical analysis was performed using an unpaired t-test (**** p < 0.0001). Scale bar is 50 μm. E, (Left panel) RPA32 (red) and gamma-H2AX (green) IF was performed in Flag-MYCN overexpressed (Dox+, from day 5 onwards) SN stage cells after DMSO or STM2457 (10 μM) treatment. STM2457 or DMSO was added from day 13 of differentiation. (Right panel) Bar plots show either RPA32 or gamma-H2AX signal intensity. Data are presented as mean ± SD from three independent experiments. Statistical analysis was performed using an unpaired t-test (** p < 0.01). Scale bar is 10 μm. F, RPA32 (red) IF was performed in SK-N-BE(2) cells with Control (shCtrl) or METTL3 KD (shMETTL3) after 48 h Dox induction. (Right panel) Bar plot shows RPA32 signal intensity. Data are presented as mean ± SD from three independent experiments. Statistical analysis was performed using an unpaired t-test (**** p < 0.0001). Scale bar is 5 μm. G, (Left panel) Representative IF showing expression of RPA32 (red) in SK-N-BE(2) cells treated either with DMSO, STM2457 (10 μM), doxorubicin (1 μM), or a combination of STM2457 with doxorubicin for 24 h. (Middle panel) Bar plot showing RPA32 signal intensity. Signal intensity measurements were taken from over 700 cells. (Right panel) Bar plot showing relative cell viability in SK-N-BE(2) cells treated for 72 h with DMSO, STM2457, Doxorubicin, and a combination of STM2457 with doxorubicin. Data are presented as mean ± SD from three independent experiments. Statistical analysis was conducted using a one-way ANOVA with Tukey’s post hoc test (*** p < 0.001, **** p < 0.0001). Scale bar is 100 μm. H, COG-N-415x patient-derived xenograft (PDX) cells were injected into NSG mice. Once tumours reached 170 mm3 mice were allocated into four treatment groups (n = 4-6 per group) and treated for 14 days with vehicle (20% hydroxypropyl-beta cyclodextrin) daily, STM2457 (50 mg/kg in vehicle) daily, doxorubicin (0.2 mg/kg in vehicle) every three days or a combination of STM2457 and doxorubicin at the same doses. Tumour volume (left panel) and body weight (right panel) is presented. Data are presented as mean ± SEM. Statistical analysis was conducted using a two-way ANOVA with Tukey’s post hoc test (*** p < 0.001, **** p < 0.0001, ns - non-significant).

To further assess the effect of STM2457 on differentiation, MYCN-amplified NB cells (SK-N-BE(2) and NGP) were treated with STM2457 in combination with RA, leading to an increase in differentiation over RA treatment alone (**Fig. 5D**; Supplementary Fig. S5B). Since HOXC9 expression was already low at the SN stage compared to tNCC (Supplementary Fig. S5C), We wanted to know what additional factors at this stage contributed to the differentiation phenotype we observed following METTL3 inhibition in MYCN overexpressed cells (**Fig. 5C**). We performed gene expression analysis of MYCN overexpressing SN stage cells treated with DMSO or STM2457. The gene expression profiles show that genes related to DNA damage and repair were upregulated during differentiation following METTL3 inhibition (Supplementary Fig. S5D). We also profiled gene expression of the METTL3 KD SK-N-BE(2) cells following RA mediated differentiation for 5 days and we observed upregulation of DNA repair related pathways in this system (Supplementary Fig. S5E). METTL3 has been shown to regulate DNA double strand break repair (2), so we reasoned that METLL3 inhibition would result in the accumulation of DNA damage because of compromised DNA repair. Indeed, MYCN overexpressing, STM2457 treated cells at the SN stage show an accumulation of the DNA damage markers RPA32 and gamma-H2AX (**Fig. 5E**). An increase in DNA damage was also detected in MYCN overexpressing, METTL3 KD cells (Supplementary Fig. S5F). We also performed RPA32 IF in METTL3 KD SK-N-BE(2) cells and again these cells showed an accumulation of DNA damage (**Fig. 5F**). MYCN overexpression creates transcriptional and replication stress thereby promoting DNA damage (50). These DNA damages need to be repaired efficiently for the survival of MYCN-amplified NB cells (51). As METTL3 inhibition enhances DNA damage in MYCN overexpressing cells, our data suggest that this accumulating DNA damage may drive proliferating MYCN overexpressing cells to differentiate. DNA-damage dependent differentiation has been observed in Leukemic cells and creating exogenous double-strand breaks by restriction enzyme was sufficient to induce differentiation (52). Consistent with this double-strand break repair related pathways were identified as top deregulated pathways in both STM2457 treated MYCN overexpressed SN stage cells and in METTL3 KD RA treated SK-N-BE(2) cells (Supplementary Fig. S5D and S5E).

We further explored METTL3 inhibition induced DNA damage as a possible combination therapy against NB. We observed that the METTL3 inhibitor STM2457 enhanced the activity of the DNA intercalating anthracycline doxorubicin in MYCN-amplified SK-N-BE(2) cells **(Fig. 5G)**. MYCN-amplified NB cells treated with a combination of STM2457, and doxorubicin accumulated higher levels of DNA damage as indicated by enhanced RPA32 IF (Fig. 5G). We next tested this combination of drugs in a patient-derived xenograft (PDX) cell line, and we found that these drugs acted synergistically resulting in reduced cell viability (Supplementary Fig. S5G). Furthermore, using an in vivo approach wherein the NSG mice with tumors resultant of subcutaneous injection of MYCN-amplified PDX cell line was treated with either vehicle control, a combination of STM2457 and doxorubicin or single drugs. We found that STM2457 along with doxorubicin was more potent in reducing tumor volume than single drugs (Fig. 5H). We also observed that none of the drugs or the combination had any significant effect on the mouse body weight (Fig. 5H). Overall, these data suggest that METTL3 inhibitors may represent efficacious therapeutic agents in the treatment of NB.

## Discussion

Our study suggests that oncogenic gene regulation by MYCN is controlled by m^6^A epitranscriptomic modification along with its well-studied role in the regulation of transcriptional process (53). We provide evidence that MYCN and METTL3 co-occupy promoter regions of the m^6^A modified genes. We provide mechanistic insight into how MYCN interaction with the m^6^A writer complex could bring m^6^A modification in developmentally regulated genes such as *HOXC8* and *HOXC9*. A similar mechanism has also been described in the case of SMAD2 which interacts with METTL3 to regulate m^6^A deposition in mRNA (6). However, sustained MYCN overexpression, as observed in MYCN-amplified NB tumors, results in aberrant epitranscriptomic regulation and deregulation of critical genes such as *HOXC8* and *HOXC9*. We show that METTL3 KD or inhibition could promote differentiation in MYCN overexpressed tNCC and MYCN-amplified NB cells (**Fig. 6**). We need further understanding of how METTL3 recruitment in gene promoter could guide m^6^A methyltransferase complex to the 3′end of the transcript which is the predominant m^6^A site identified in mature transcript (8). We observed m^6^A positive genes in NB tumors and MYCN/METTL3 co-bound genes which were positive of m^6^A in tNCC were related to axonogenesis (**Fig. 2D**; Supplementary Figure S4D). This suggests, MYCN and METTL3 mediated epitranscriptomic regulation might play a wider role apart from HOX gene regulation and could be a key player in MYCN induced oncogenic transformation of the tNCC (12,14), which required further investigation. Our study evidence paves the way for further studies on the mechanisms of m^6^A epigenome modification through MYCN or other oncogenes, in NB and other cancers, by guiding METTL3 to specific genomic locus.

**Figure 6.**
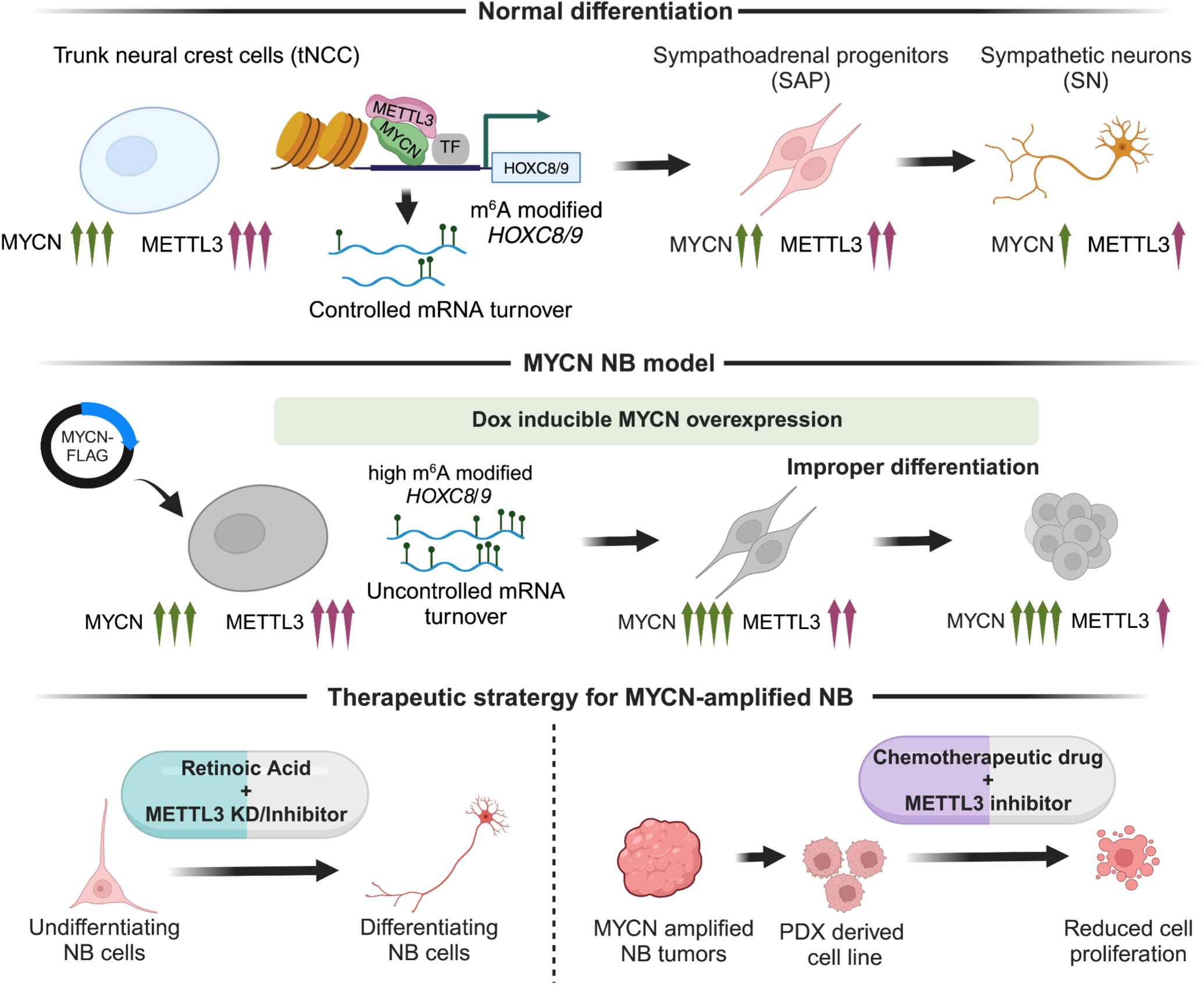
The model illustrates (top panel) the dynamic fine-tuning of developmentally important posterior HOX genes, during differentiation of tNCC and further to SN by cooperation of MYCN and METTL3. METTL3 deposits m6A RNA modification on HOX genes, thereby facilitating controlled mRNA turnover leading to a normal differentiation process. (middle panel) To better comprehend the role of METTL3 and MYCN during early differentiation, we created an MYCN NB model by overexpressing MYCN during the tNCC-SN differentiation process. The MYCN overexpression led to an increase in m6A modification of HOX genes and further the tNCC failed to differentiate to SN (bottom panel) Finally, we designed two therapeutic strategies using METTL3 inhibitor to treat MYCN amplified high-risk NB tumors. As the NB cells failed to differentiate, we utilized RA along with METTL3 inhibitor (STM2457) and observed restoration of differentiation phenotype. Combining doxorubicin and STM2457 had a synergistic effect on cell viability.

HOXC9 gene was identified before as the top significantly downregulated genes in high-risk NB (43). A study on epigenetic regulation, such as analysis of DNA methylation could not explain the downregulation of *HOXC9* gene expression in high-risk NB (43). We uncover the unexpected role of METTL3 mediated m^6^A modification in controlling HOX gene expression in NB. We provide evidence that m^6^A epitranscriptomic modification could explain deregulation in HOX gene expression in NB. Differentiation of tNCC needs to be regulated tightly but also dynamically. The tNCC are on the move following delamination and needs to differentiate at different time points throughout their migration during development. We propose that the m^6^A epitranscriptome mediated gene regulation provides flexibility by rapidly regulating important lineage specific transcription factors as and when required during the differentiation of the crest cells. Epigenetic regulation of HOX gene is well studied (54) but we provide evidence on a m^6^A epitranscriptomic modification-dependent regulation of the HOX genes for the first time in NB. Several studies have implicated the phenotypic plasticity of NB tumor cells, Chromatin modification dependent epigenetic mechanisms along with external environmental cues have been implicated in such phenotypic plasticity (55,56). Given that m^6^A modification can regulate epithelial to mesenchymal transition (57), further studies are required to reveal if epitranscriptomic-based regulation contributes to phenotypic plasticity in NB tumors.

Apart from its role in the regulation of post-transcriptional gene expression such as RNA stability, recent studies have shown a role of m^6^A modification in gene transcription (58). METTL3 mediated m^6^A modification of promoter associated RNA can recruit polycomb repressive complex 2 in YTHDC1 dependent manner (7). Our data on promoter bound METTL3 suggest that m^6^A modification can control epigenetic state in differentiating tNCC and this might contribute to a widespread deregulation in gene expression observed following METTL3 KD. We speculate METTL3 and MYCN co-binding at the gene promoter can drive m^6^A modification of promoter-associated transcripts, thereby could affect the epigenetic state of NB cells which require further investigation.

Our gene expression data suggest genes related to DNA damage response were upregulated when METTL3 was inhibited using small molecule in MYCN overexpressed SN stage cells. Consistent with this we could detect the increase in DNA damage markers following METTL3 KD and inhibition in MYCN overexpressed SN stage cells. We believe induction of DNA damage following pharmacologic inhibition of METTL3 acts as a further trigger for differentiation of the MYCN overexpressed cells apart from *HOXC8*/*HOXC9* upregulation. Differentiation induced by DNA damage has been reported before in several other experimental models (52,59). We explored this further therapeutically in MYCN-amplified NB cells and PDX cell lines where METTL3 pharmacological inhibition combined with doxorubicin was more effective in inhibiting cell viability, compared to the single agents. We propose treating with METTL3 inhibitor to sensitize the tumor cells to chemotherapeutic drugs could be an effective treatment strategy. There is a growing interest in developing more effective METTL3 inhibitors and recently STM3006 was described as 20 times more potent than STM2457 in cell-based assays. However, the bioavailability of STM3006 was limited, because it was rapidly metabolized, highly reducing the drug effectiveness *in vivo* (60).

Overall, our findings reveal that MYCN can cooperate with METTL3 to establish an m^6^A epitranscriptomic signature over developmentally regulated *HOXC8* and *HOXC9* genes. We provide evidence that pharmacological inhibition of METTL3 could be a novel therapeutic approach for high-risk NB, by inducing differentiation and increasing the efficacy of the chemotherapeutic drugs (**Fig. 6**).

## Data availability

The data supporting the findings of this article are accessible through the NCBI Gene Expression Omnibus (GEO) at https://www.ncbi.nlm.nih.gov/geo/. These data are associated with the GSE244473 accession number.

## Acknowledgments

We would like to thank the core facility at Novum, BEA, Bioinformatics and Expression Analysis, which is supported by the board of research at the Karolinska Institute and the research committee at the Karolinska Hospital for help with sequencing. The computations and data handling were enabled by resources in project SNIC-2022-22-85 provided by the Swedish National Infrastructure for Computing (SNIC) at UPPMAX, partially funded by the Swedish Council through grant agreement no. 2018-05973.

