## Supplementary data for "METTL3/MYCN cooperation drives neural crest differentiation and provides therapeutic vulnerability in neuroblastoma"

Thombare, Vaid, et al

Department of Laboratory Medicine, Institute of Biomedicine, University of Gothenburg, Gothenburg, Sweden

**Contents**

Supplementary Figures S1-S5.

Supplementary method

Reference to the supplementary data

**Other Supplementary Materials**

Supplementary Table S1: Sequence of siRNAs, shRNAs, qPCR primers, plasmids, and reagents used in the study. Also contains information regarding all the antibodies used in the study.

Supplementary Table S2: List of all the samples sequenced.

#### Supplementary Figure S1

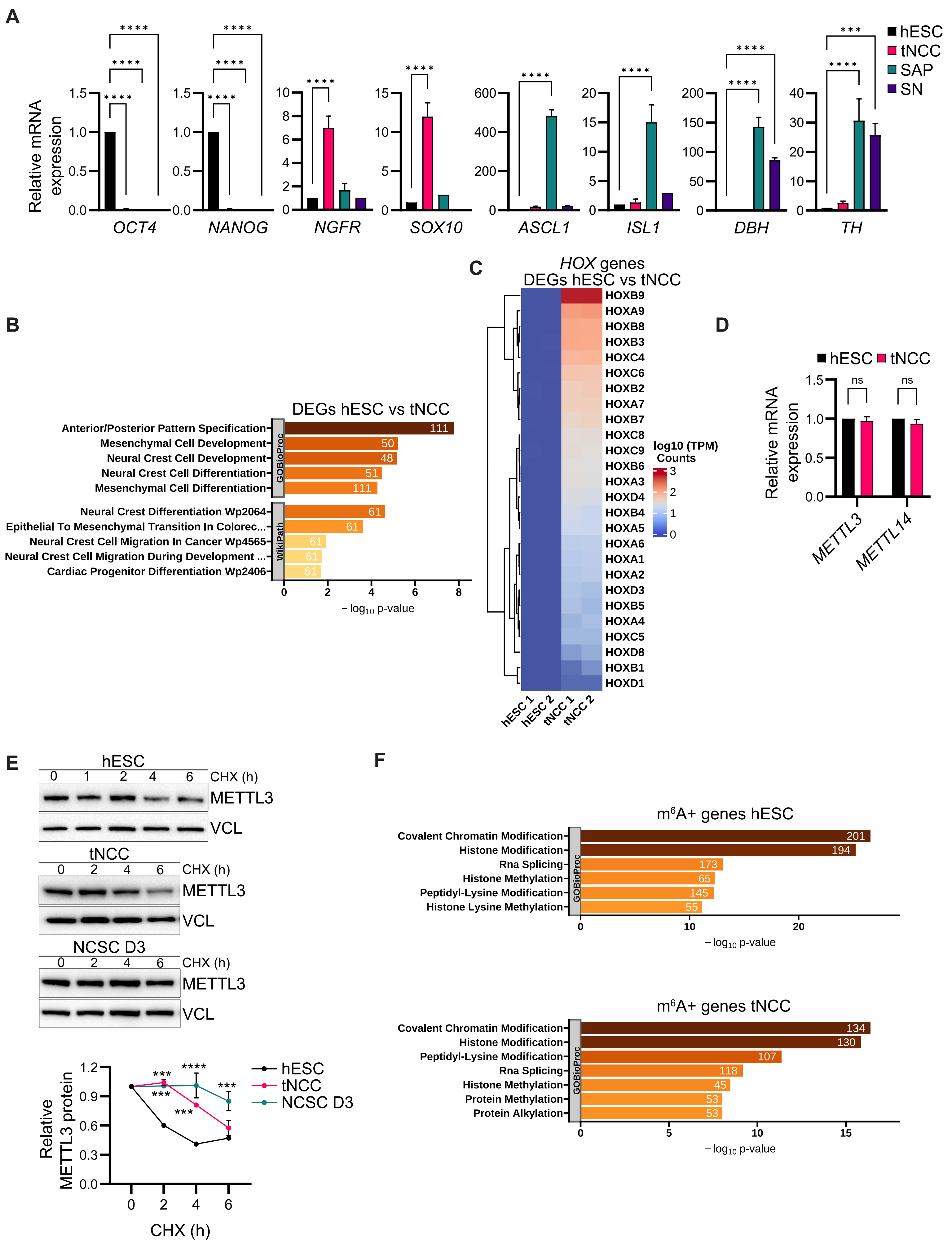

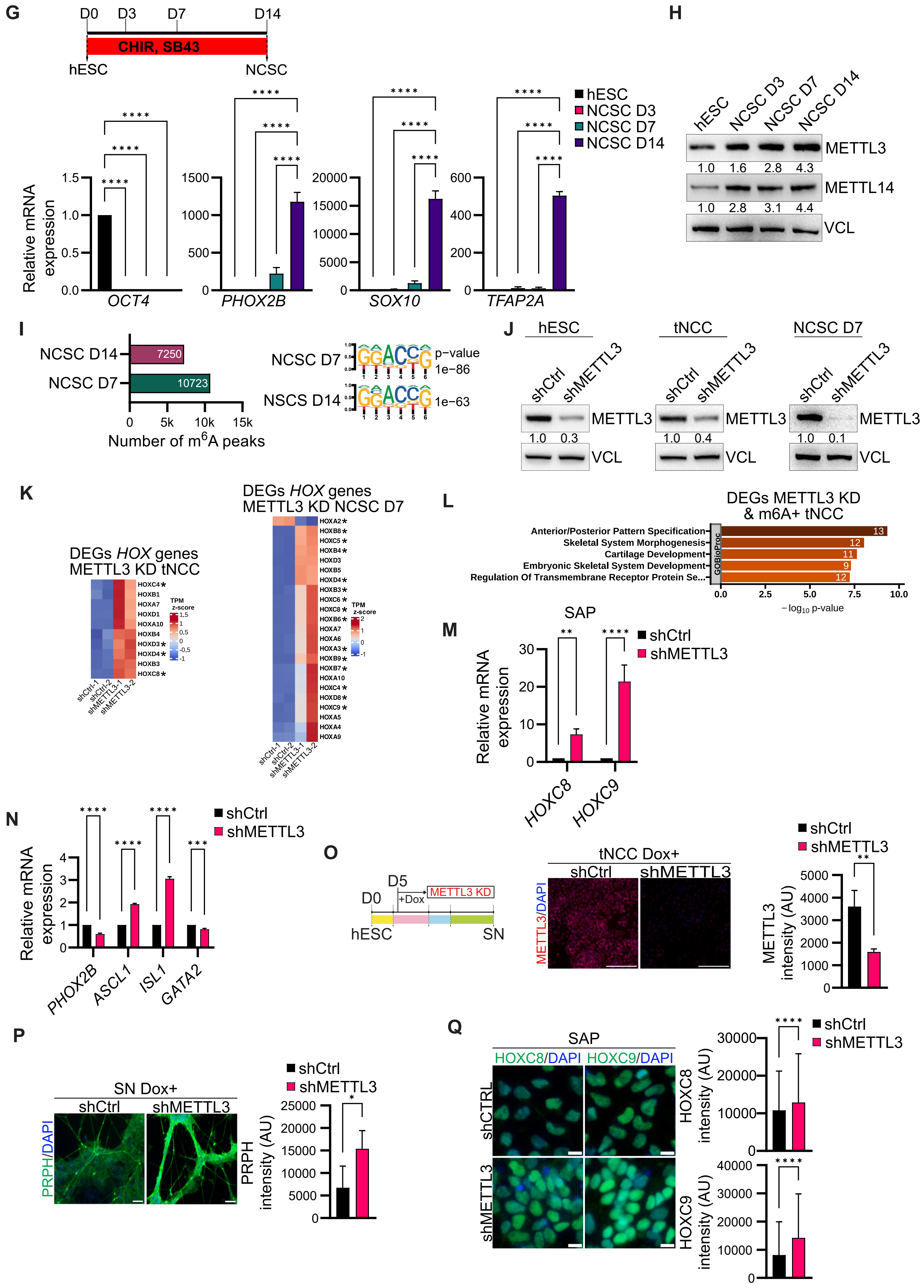

G
D0 D3 D7 D14
CHIR, SB43
hESC NCSC
Relative mRNA expression
OCT4
PHOX2B
SOX10
TFAP2A
hESC
NCSC D3
NCSC D7
NCSC D14
H
hESC NCSC D3 NCSC D7 NCSC D14
METTL3
1.0 1.6 2.8 4.3
METTL14
1.0 2.8 3.1 4.4
VCL
I
NCSC D14 7250
NCSC D7 10723
Number of m6A peaks
NCSC D7 p-value 1e-86
NSCS D14 1e-63
J
hESC tNCC NCSC D7
shCtrl shMETTL3
METTL3
1.0 0.3
1.0 0.4
1.0 0.1
VCL
K
DEGs HOX genes METTL3 KD tNCC
DEGs HOX genes METTL3 KD NCSC D7
TPM z-score
L
DEGs METTL3 KD & m6A+ tNCC
Anterior/Posterior Pattern Specification 13
Skeletal System Morphogenesis 12
Cartilage Development 11
Embryonic Skeletal System Development 9
Regulation Of Transmembrane Receptor Protein Se... 12
GOBioProc
−log10 p-value
M
SAP
shCtrl
shMETTL3
HOXC8 HOXC9
N
PHOX2B ASCL1 ISL1 GATA2
O
D0 D5 +Dox METTL3 KD
hESC SN
tNCC Dox+
METTL3/DAPI
METTL3 intensity (AU)
P
SN Dox+
PRPH/DAPI
PRPH intensity (AU)
Q
HOXC8/DAPI HOXC9/DAPI
shCTRL
shMETTL3
HOXC8 intensity (AU)
HOXC9 intensity (AU)

**Supplementary Figure S1. A,** Bar plots depict the relative mRNA expression levels of lineage markers at different stages, including hESC, tNCC, SAP, and SN. Pluripotency markers NANOG and OCT4, tNCC markers NGFR and SOX10, SAP markers ASCL1 and ISL1 and SAP/SN markers DBH and TH were quantified. GAPDH served as a reference for qPCR data normalization. The presented data represents the mean ± SD from three independent biological replicates. Statistical analysis was conducted using a two-way ANOVA with Tukey's *post hoc* test (**** $p< 0.0001$). **B,** Top enriched terms associated with DEGs (hESC vs. tNCC). **C,** Heatmap demonstrating the top differentially expressed HOX genes between hESC and tNCC. **D,** Relative mRNA expression of METTL3 and METTL14 in hESC and tNCC as determined by RT-qPCR. GAPDH served as the normalization reference for qPCR data. The results are presented as mean ± SD from three independent biological replicates. Statistical analysis was conducted using two-way ANOVA with Šídák's multiple comparisons test, and non-significant differences (ns) indicate $p > 0.05$. **E,** In the top panel, an immunoblot displays the levels of METTL3 in hESC, tNCC, and neural crest stem cells (NCSC) at day 3 following cycloheximide (CHX) chase for specified time intervals. Vinculin was utilized as the loading control. The bottom panel illustrates line plots presenting the quantification of METTL3 levels after CHX chase at the indicated time points. The experiments were replicated three times, and the data are presented as mean ± SD in the graph. Statistical analysis was performed using two-way ANOVA with Dunnett's multiple comparisons test (*** $p < 0.001$, **** $p < 0.0001$). **F,** Top enriched terms associated with the $m^6A$ positive ($m^6A$ +) genes in hESC (top panel) and tNCC (bottom panel). **G,** The schematic diagram delineates the critical stages in the differentiation of hESC into neural crest stem cells (NCSC). In the bottom panel, the relative mRNA expression levels of lineage markers at the hESC and NCSC stages are presented. OCT4 serves as a pluripotency marker, while PHOX2B, SOX10, and TFAP2A are employed as markers for NCSC. GAPDH is utilized for normalizing the qPCR data. The data is represented as mean ± SD of three replicates, and statistical analysis was conducted using two-way ANOVA with Tukey's post hoc test (**** $p < 0.0001$). **H,** Representative immunoblot shows the expression levels of METTL3 and METTL14 in hESC and at various stages during NCSC differentiation. Vinculin serves as the loading control, and the values below the blots indicate the fold change in the levels of METTL3 and METTL14. **I,** The left panel displays the total number of $m^6A$ peaks in day 7 NCSC progenitors and day 14 NCSC, while the right panel presents the motifs identified through de novo motif analysis in the $m^6A$ peaks. **J,** Immunoblot shows METTL3 KD in hESC (left panel), tNCC (middle panel), and NCSC day 7 (right panel). Vinculin was used as a loading control. The values below indicate the fold change in levels of METTL3. **K,** Heatmap summarizes the top differentially expressed HOX genes between control and METTL3 KD tNCC and in NCSC. The '*' indicates the presence of $m^6A$ peaks in the gene. **L,** Top enriched terms associated with DEGs (shCtrl vs. METTL3 KD) having $m^6A$ peaks ($m^6A$+) in tNCC. **M,** RT-qPCR data showing the expression of *HOXC8* and *HOXC9* in SAP following METTL3 KD. GAPDH was used to normalize the qPCR data. Data are shown as mean ± SD of three replicates. Two-way ANOVA with Šídák's multiple comparisons test was used (** $p < 0.01$, **** $p < 0.0001$). **N,** RT-qPCR data showing the expression of SAP markers *PHOX2B*, *ASCL1*, *ISL1*, and *GATA2* after METTL3 KD. *GAPDH* was used to normalize the qPCR data. Data are presented as mean ± SD of three replicates. Two-way ANOVA with Šídák's multiple comparisons test was employed (*** $p < 0.001$, **** $p < 0.0001$). **O,** In the left panel, a schematic diagram provides a timeline of METTL3 KD in Dox dependent manner during tNCC differentiation. The middle panel displays representative immunofluorescence (IF) images showing METTL3 (red) expression in tNCC, confirming METTL3 KD following Dox induction. The right panel shows the quantification of METTL3 signal intensity after METTL3 KD. Statistical analysis was performed using an unpaired *t*-test (** $p < 0.01$). Scale bar represents 100 µm. **P,** Dox induced shCtrl and METTL3 KD (shMETTL3) tNCC were differentiated to SN and IF was performed with PRPH (green) antibody. The Bar plot shows PRPH signal intensity. Experiments were performed in three independent biological replicates. Unpaired *t*-test was used, * $p < 0.05$. Scale bar represents 100 µm. **Q,** Dox induced shCtrl and METTL3 KD (shMETTL3) tNCC were differentiated to SAP and IF was performed with HOXC8 and HOXC9 antibodies. Bar plots illustrate the quantification of signal intensity for HOXC8 and HOXC9. Data is presented as mean ± SD and three independent experiments were conducted, with signal intensity measurements taken from over 1000 cells. Scale bar represents 10 µm. Statistical analysis was performed using an unpaired *t*-test (**** $p < 0.0001$).

### Supplementary Figure S2

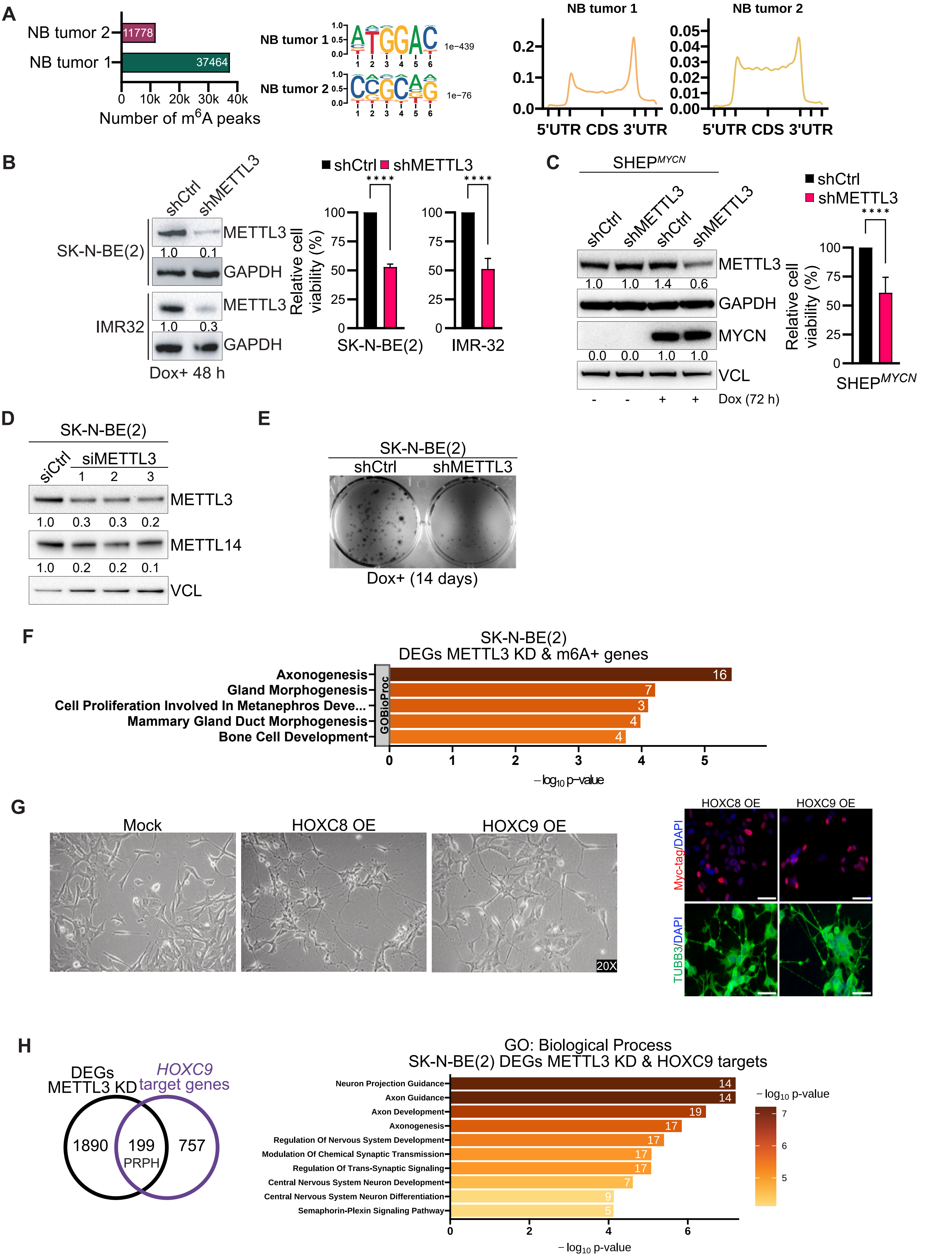

I

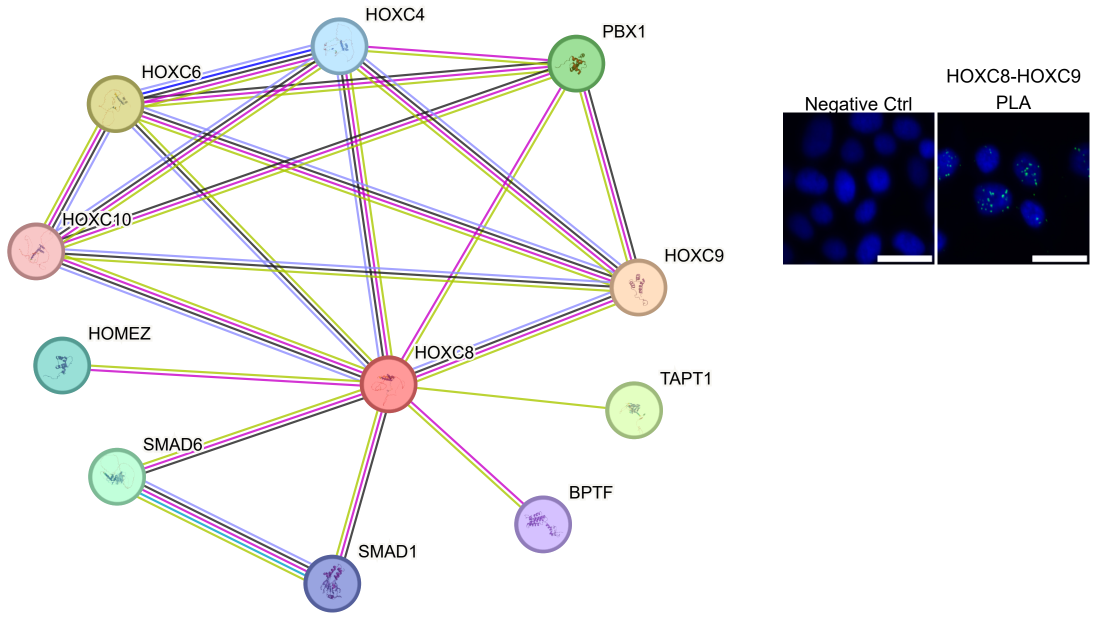

HOXC4
PBX1
HOXC6
HOXC10
HOXC9
HOMEZ
HOXC8
TAPT1
SMAD6
BPTF
SMAD1
Negative Ctrl
HOXC8-HOXC9
PLA


J

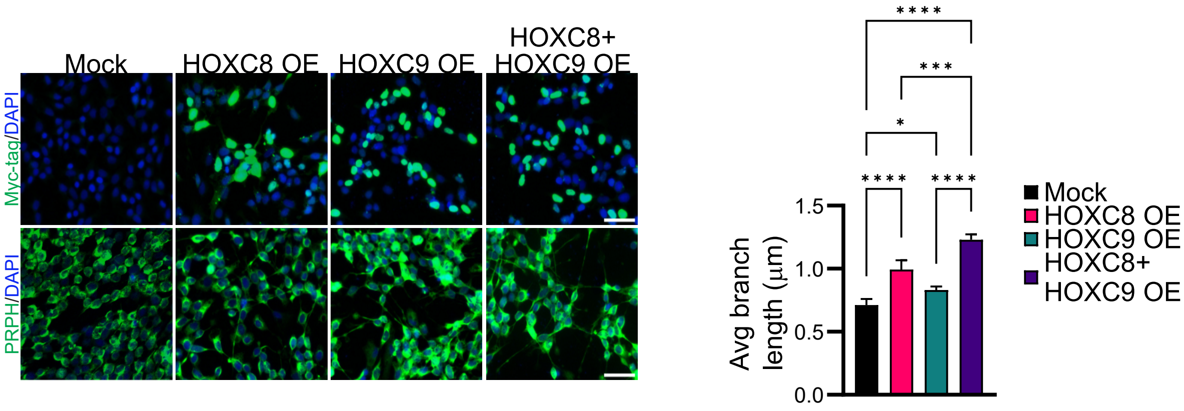

Mock
HOXC8 OE
HOXC9 OE
HOXC8+
HOXC9 OE
Myc-tag/DAPI
PRPH/DAPI
****
***
*
****
****
1.5
1.0
0.5
0.0
Avg branch
length (μm)
Mock
HOXC8 OE
HOXC9 OE
HOXC8+
HOXC9 OE

**Supplementary Figure S2. A,** (Left panel) The total number of m$^6$A peaks identified and (middle panel) identified motifs from de novo motif analysis of m$^6$A peaks enriched in MYCN-amplified NB tumor samples. (Right panel) Metagene analysis showing relative m$^6$A peak density at genes in both MYCN-amplified NB tumor samples. **B,** (Left panel) Immunoblot showing METTL3 expression in Dox induced (48 hours) control and METTL3 KD SK-N-BE(2) and IMR-32 cells . GAPDH was used as a loading control. The values below indicate the fold change in levels of METTL3. (Right panel) Bar plots show cell viability of SK-N-BE(2) and IMR-32 cells with METTL3 KD (6 days post Dox induction). Data are presented as mean ± SD from three independent experiments. Unpaired *t*-test was used, **** $p < 0.0001$. **C,** (Left panel) Representative immunoblot showing simultaneous MYCN overexpression and METTL3 KD *via* Dox induction in SHEP (SHEP$^{MYCN}$) cells. GAPDH and vinculin were loading controls. The values below indicate the fold change in levels of METTL3 and MYCN. (Right panel) Bar plot shows the cell viability of SHEP$^{MYCN}$ cells with METTL3 KD (6 days post Dox induction). Data are presented as mean ± SD from three independent experiments. Unpaired *t*-test was used, **** $p < 0.0001$. **D,** Representative immunoblot showing METTL3 and METTL14 expression following siRNA-mediated METTL3 KD. Vinculin was used as a loading control. The values below indicate the fold change in levels of METTL3 and METTL14. **E,** Representative images from colony formation assay performed in SK-N-BE(2) cells with METTL3 KD (14 days post Dox induction). **F,** Top enriched terms associated with DEGs (shCtrl Vs. METTL3 KD) having m$^6$A peaks (m$^6$A+) in SK-N-BE(2) cells. **G,** (Left panel) Brightfield images of SK-N-BE(2) cells with stable overexpression of HOXC8 and HOXC9. (Right panel) Representative IF showing TUBB3 (green) and overexpression of MYC-tagged HOXC8 and HOXC9 (red) in SK-N- BE(2) cells. **H,** (Left panel) Venn diagram comparison of HOXC9 target genes (1) [genes with HOXC9 ChIP-seq peak and 1.5 fold change in expression between control vs HOXC9 overexpression] and differentially expressed between DEGs (shCtrl Vs. METTL3 KD) in SK-N-BE(2) cells. (Right panel) Top enriched terms associated with the overlapping genes from the left panel. **I,** (Left panel) Interaction network of HOXC8 obtained using STRING-db with default parameters. (Right panel) Proximity ligation assay (PLA) showing the HOXC8 and HOXC9 PLA signal (green) between in SK-N-BE(2) cell nucleus (marked by DAPI). The Negative control shows PLA with only the HOXC8 antibody. Scale bar represents 50 µm. **J,** (Left panel) Representative IF showing staining for PRPH and MYC- tagged HOXC8 and HOXC9 in SK-N-BE(2) cell transiently overexpressing HOXC8 and HOXC9 individually or in combination followed by retinoic acid (RA) mediated differentiation for 3 days. (Right panel) Bar graph shows the quantification of the average neurite branch length. Scale bar represents 50 µm. Data are presented as mean ± SD from three independent experiments. Two-way ANOVA with Tukey's *post hoc* test was used, * $p < 0.05$, *** $p < 0.001$, **** $p < 0.0001$.

#### Supplementary Figure S3

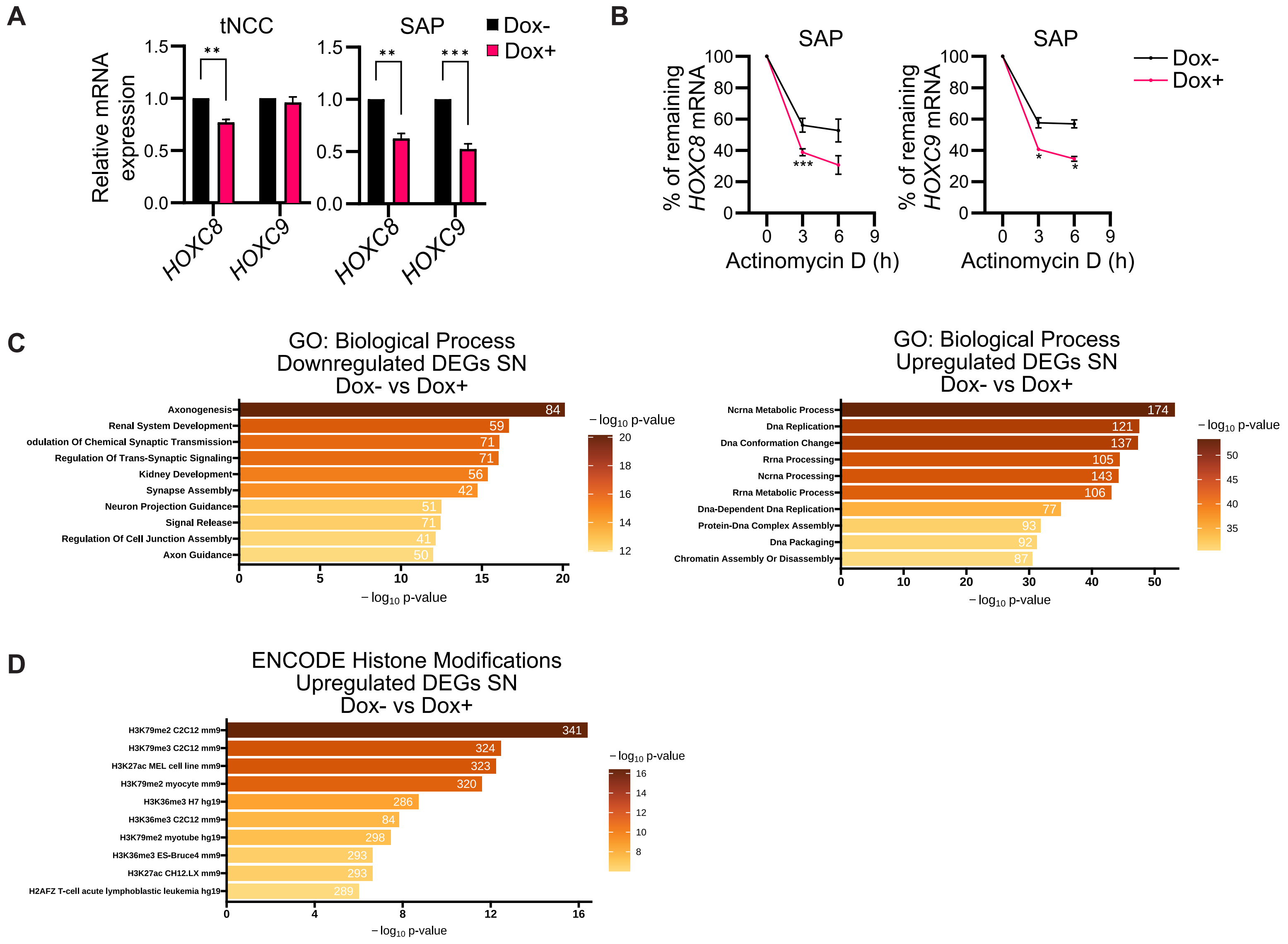

**Supplementary Figure S3. A,** Relative mRNA expression of *HOXC8* and *HOXC9* in tNCC and SAP following Flag- MYCN overexpression (Dox +, from day 5 onwards) and in control (Dox-). GAPDH was used to normalize the qPCR data. Data are shown as mean ± SD of three replicates. Statistical analysis was performed using an unpaired *t*-test, ** $p < 0.01$, *** $p < 0.001$. **B,** Stability of *HOXC8* and *HOXC9* transcripts detected by RT-qPCR after Actinomycin D (10 μg/ml) mediated transcription blocking for the time points indicated in MYCN overexpressed SAP. Line plots presenting the quantification of remaining levels of *HOXC8* and *HOXC9* transcript at the indicated time points (n=3). Two-way ANOVA with Šídák's multiple comparisons test was employed (* $p < 0.05$, *** $p < 0.001$). **C,** Top enriched terms associated with downregulated (left) and upregulated (right) genes in Control (Dox-) vs. Flag-MYCN overexpressed (Dox +, from day 5 onwards) SN stage cells. **D,** Top enriched terms associated with histone modificatioins in upregulated geneset obtained from Control (Dox-) vs. Flag-MYCN overexpressed (Dox +, from day 5 onwards) SN stage cells.

#### Supplementary Figure S4

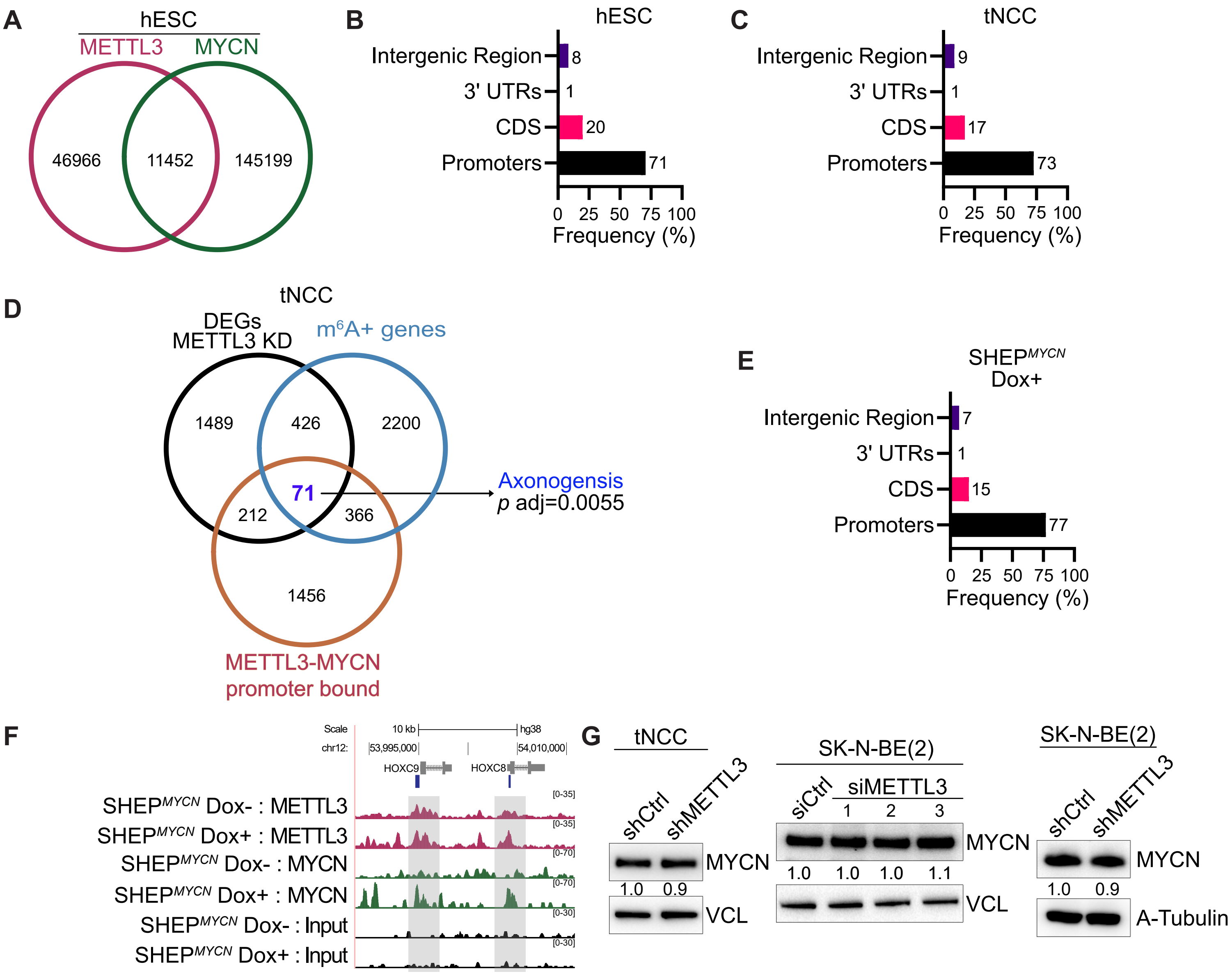

**Supplementary Figure S4. A,** Venn diagram comparison of METTL3 and MYCN binding sites determined from ChIP-seq experiments performed in hESC. **B, C,** Distribution of METTL3 and MYCN co-bound regions by genomic features for (B) hESC and (C) tNCC. METTL3 and MYCN co-bound regions were determined using the ChIP-seq experiments. **D,** Three-way Venn diagram comparing DEGs (shCtrl vs. shMETTL3), $m^6A$ positive genes, and METTL3-MYCN promoter bound regions in tNCC. Top enriched term associated with 71 genes that were common in all three conditions is highlighted. **E,** Distribution of METTL3 and MYCN co-bound regions by genomic features for SHEP$^{MYCN}$ cells after Dox induction. METTL3 and MYCN co-bound regions were determined using the ChIP-seq experiments. **F,** Genome Browser screenshot showing METTL3, MYCN ChIP-seq signals over the HOXC8 and HOXC9 gene locus in SHEP$^{MYCN}$ cells before and after Dox induction. METTL3 and MYCN overlapping peak coordinates in SHEP$^{MYCN}$ after Dox induction are indicated by blue bars. **G,** Immunoblot showing MYCN expression following METTL3 KD in tNCC, SK-N-BE(2) cells. The values below indicate the fold change in levels of MYCN. METTL3 KD blots are presented in Supplementary Fig. S1J, S2D, and S2B respectively.

#### Supplementary Figure S5

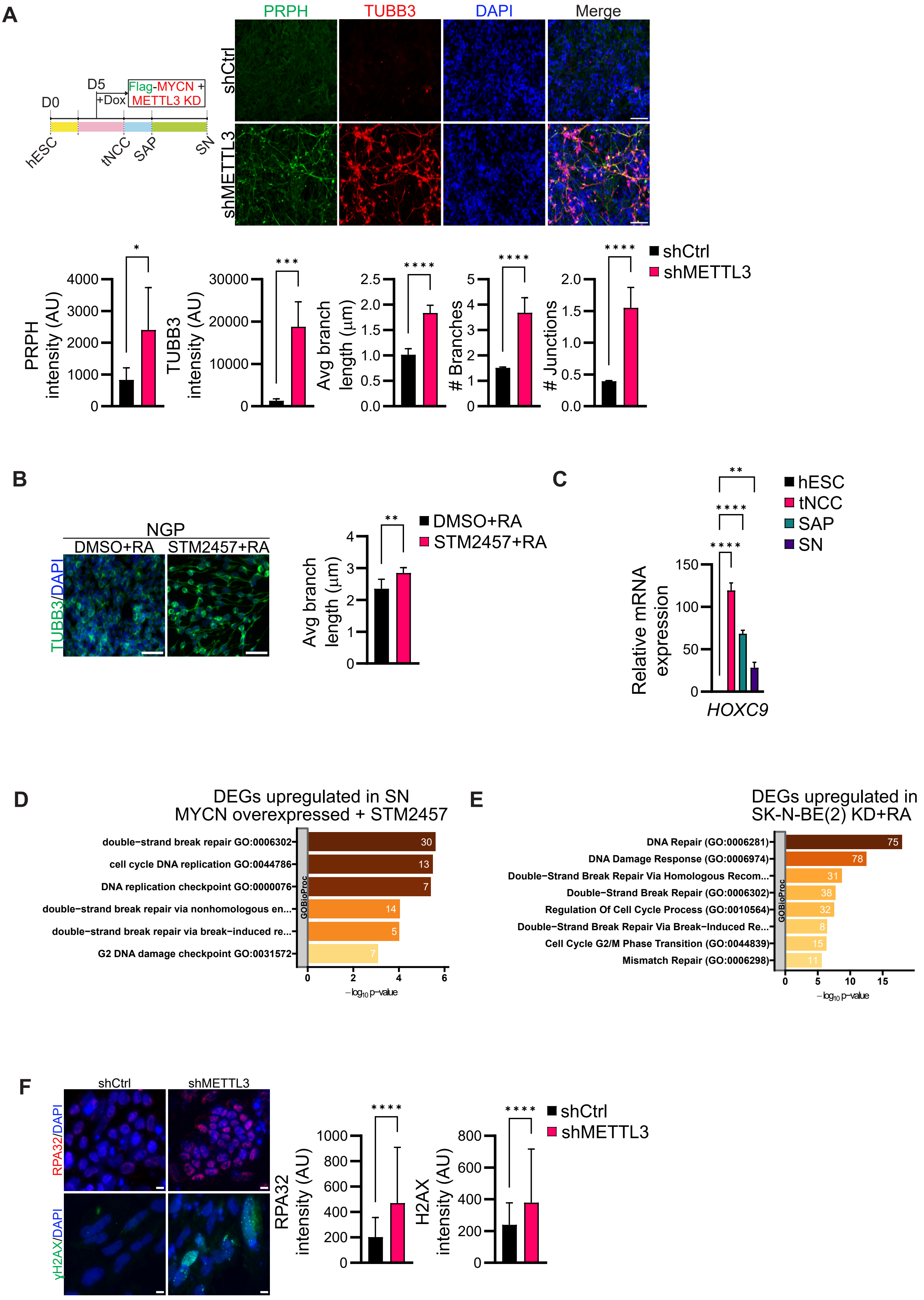

G

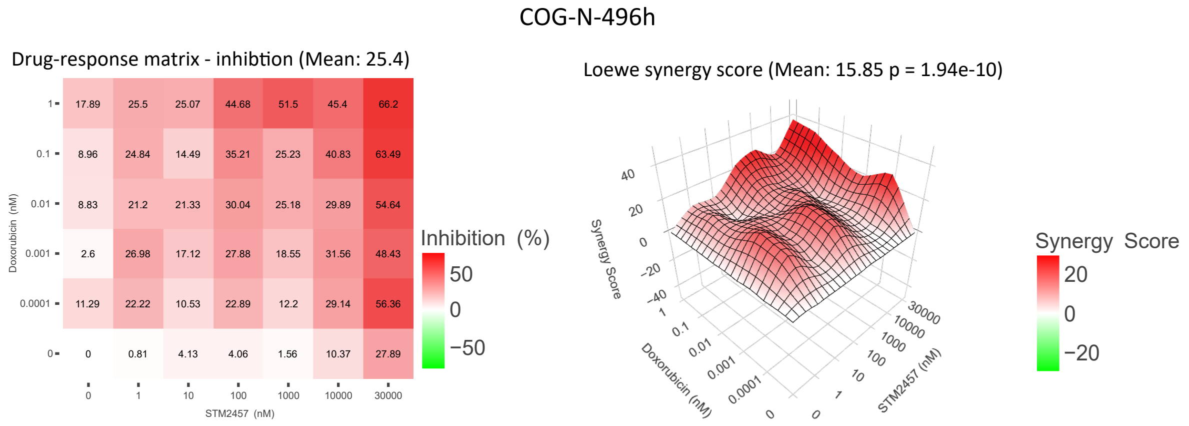
COG-N-496h
Drug-response matrix - inhibtion (Mean: 25.4)
1 17.89 25.5 25.07 44.68 51.5 45.4 66.2
0.1 8.96 24.84 14.49 35.21 25.23 40.83 63.49
0.01 8.83 21.2 21.33 30.04 25.18 29.89 54.64
0.001 2.6 26.98 17.12 27.88 18.55 31.56 48.43
0.0001 11.29 22.22 10.53 22.89 12.2 29.14 56.36
0 0 0.81 4.13 4.06 1.56 10.37 27.89
0 1 10 100 1000 10000 30000
STM2457 (nM)
Doxorubicin (nM)
Inhibition (%)
50
0
−50
Loewe synergy score (Mean: 15.85 p = 1.94e-10)
Synergy Score
40
20
0
−20
−40
Doxorubicin (nM)
1
0.1
0.01
0.001
0.0001
0
STM2457 (nM)
30000
10000
1000
100
10
1
0
Synergy Score
20
0
−20

**Supplementary Figure S5. A,** (Left panel) Schematic diagram showing a timeline of simultaneous induction of Flag- MYCN and METTL3 KD during differentiation (Dox+, from day 5 onwards). (Right panel) Representative IF showing PRPH (green) and TUBB3 (red) expression in SN stage cells with simultaneous Flag-MYCN overexpression and METTL3 KD induced by Dox at the indicated day. Bar plots showing either PRPH, TUBB3 intensity, or quantification of neurite branch length, number of branches, and junctions. Data are shown as mean ± SD and this analysis was conducted across three independent biological replicates. Statistical significance was determined using an unpaired *t*-test (* $p < 0.05$, *** $p < 0.001$, **** $p < 0.0001$). Scale bar represents 100 μm. **B,** Representative IF images of TUBB3 (green) in NGP cells that were pre-treated with either DMSO or STM2457 (10 μM) for 24 hours, followed by a RA treatment for another 3 days. (Right panel) Bar plot shows quantification of neurite branch length. Data are presented as mean ± SD from three independent experiments. Statistical significance was determined using an unpaired *t*-test (** $p < 0.01$). Scale bar represents 50 μm. **C,** Relative mRNA expression of *HOXC9* in hESC, tNCC, SAP, and SN. GAPDH was used to normalize the qPCR data. Data are shown as mean ± SD of three replicates. Two-tailed paired *t*-test was used, ** $p < 0.01$ **** $p < 0.0001$. **D,** Top enriched terms associated with upregulated genes in Flag-MYCN overexpressed (Dox+, from day 5 onwards) SN stage cells (day 20 of differentiation) after DMSO or STM2457 (10 μM) treatment. STM2457 or DMSO was added from day 13 of differentiation. **E,** Top enriched terms associated with upregulated DEGs following Dox induced METTL3 KD and RA treatment in SK-N-BE(2) cells for 5 days. **F,** (Left panel) Representative IF showing expression of RPA32 (red) and gamma H2AX (green) expression in SN stage cells with simultaneous Flag-MYCN overexpression and METTL3 KD induced by Dox as indicated in panel (A) above. (Right panel) Bar plots show either RPA32 or gamma-H2AX intensity. Data is presented as mean ± SD and three independent experiments were conducted, with signal intensity measurements taken from over 500 cells. Scale bar represents 10 μm. Statistical analysis was performed using an unpaired *t*-test (**** $p< 0.0001$). **G,** (Left panel) Dose-response matrix of STM2457 plus doxorubicin in COG-N-496h patient-derived xenograft (PDX) line, treated for 72 h with the combination of agents or agents alone; color gradients represent the % of inhibition in viability compared to the DMSO vehicle control treated cells. (Right panel) Loewe synergy scores were calculated with Synergy Finder and are shown in the ZIP score Synergy map. Scores >10 represent synergism in the activity of the drugs. Results shown are from the average of two independent experiments each with technical replicates.

**Supplementary method:**

**Chemical synthesis of STM2457**

***N*-[(6-{[(cyclohexylmethyl)amino]methyl}imidazo[1,2-a]pyridin-2-yl)methyl]-4-oxo-4*H*pyrido[1,2-a]pyrimidine-2-carboxamide – STM2457**

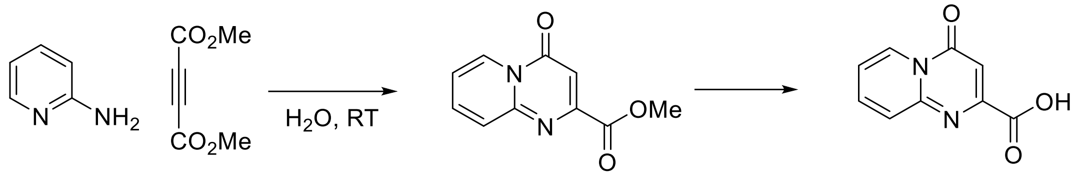


**Scheme 1. Synthesis of 4-oxo-4*H*-pyrido[1,2-a]pyrimidine-2-carboxylic acid**

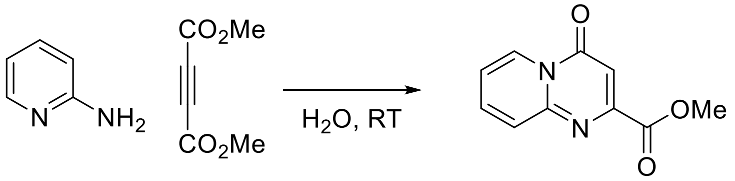


**Step 1: Methyl 4-oxopyrido[1,2-a]pyrimidine-2-carboxylate.**

Dimethyl but-2-ynedioate (16 mL, 0.126 mol) was added slowly to a vigorously stirred solution of pyridin-2-amine (10.00 g, 0.105 mol) in water (1000 mL), and the reaction mixture was stirred at room temperature for 18 hours. After completion of reaction, the reaction mixture was extracted with DCM (3x100 mL), organic layer was collected and dried over $MgSO_4$ and volatiles were removed under reduced pressure. The crude product was purified by column chromatography on silica gel using eluent ethyl acetate in hexane to obtain methyl 4-oxopyrido[1,2-a]pyrimidine-2-carboxylate (8.7 g, 40%) as an off-white solid.

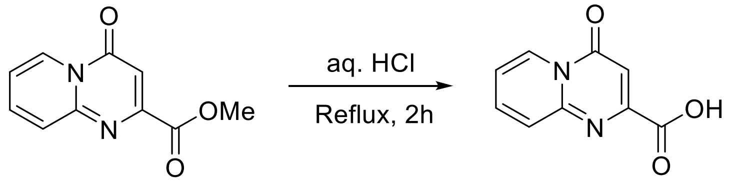


**Step 2: 4-oxo-4*H*-pyrido[1,2-a]pyrimidine-2-carboxylic acid.**

Methyl 4-oxopyrido[1,2-a]pyrimidine-2-carboxylate (8.7 g, 42.9 mmol) was dissolved in aq. HCl (~8M, 9.5 mL) at room temperature and the solution was heated at reflux for 2 hours. The mixture was cooled to room temperature and the precipitate was collected by filtration and dried on the filter to give the title compound 4-oxo-4*H*-pyrido[1,2-a]pyrimidine-2-carboxylic acid. (6.5 g, 79 %). $^{1}$H NMR (600 MHz, DMSO-$d_6$) $\delta$ 9.00 (d, J = 6.3 Hz, 1H), 8.05 (ddd, J = 8.6, 6.7, 1.6 Hz, 1H), 7.83 (dt, J = 8.8, 1.1 Hz, 1H), 7.46 (td, J = 6.9, 1.4 Hz, 1H), 6.87 (s, 1H).

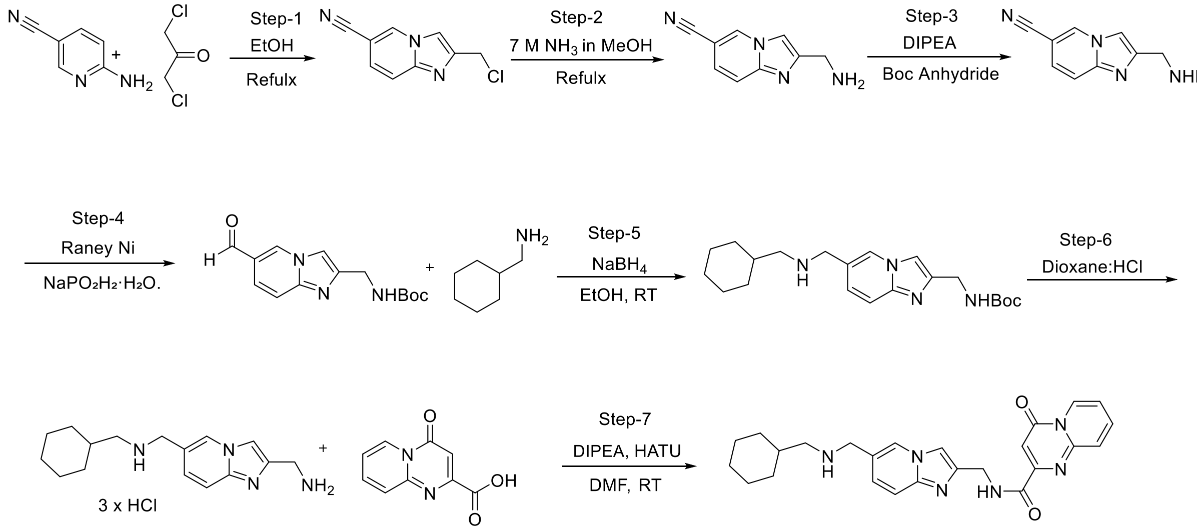


**Scheme 2: Synthesis of STM 2457**

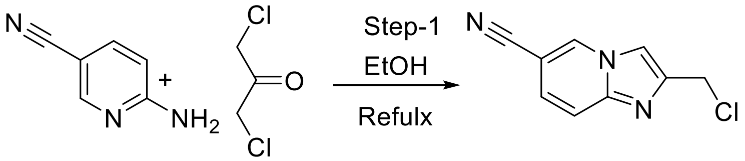


**Step 1: 2-(Chloromethyl)imidazo[1,2-*a*]pyridine-6-carbonitrile**

To a stirred solution of 6-aminonicotinonitrile (15 g, 126 mmol) in ethanol (40 mL), 1,3-dichloropropan-2-one (45g, 378 mmol) was added under nitrogen atmosphere and stirred for 16 h at 60 °C. After completion of the reaction, the reaction mixture was poured into ice cold saturated solution of sodium bicarbonate (160 mL), then filtered to obtain a crude product. The crude product was purified by column chromatography on silica gel using eluent ethyl acetate in hexane to obtain 2-(Chloromethyl)imidazo[1,2-a]pyridine-6-carbonitrile (20 g, 104 mmol, 83 % yield). $^{1}$H NMR (600 MHz, $CDCl_3$) $\delta$ 8.53 (dd, $J$ = 1.7, 1.0 Hz, 1H), 7.71 (d, $J$ = 0.8 Hz, 1H), 7.64 (dt, $J$ = 9.4, 0.9 Hz, 1H), 7.31 – 7.26 (m, 1H), 4.74 (d, $J$ = 0.7 Hz, 2H).

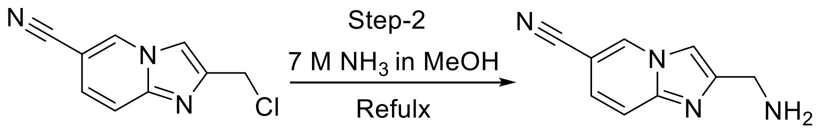


**Step 2: 2-(Aminomethyl)imidazo[1,2-*a*]pyridine-6-carbonitrile**

To a stirred solution of 2-(chloromethyl)imidazo[l,2-a]pyridine-6-carbonitrile (5 g, 26.17 mmol) in methanol (50 mL), (7N) $NH_3$ in methanol (4.47 mL, 67.6 mmol) was added and stirred for 4 h at 80 °C. The volatiles were removed under reduced pressure to obtain a 2-(aminomethyl)imidazo[1,2-a]pyridine-6-carbonitrile (4.2 g, 83 % yield).

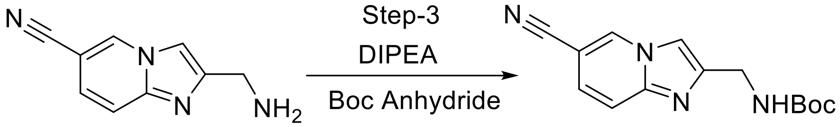


**Step 3: *tert*-Butyl ((6-cyanoimidazo[1,2-*a*]pyridin-2-yl)methyl)carbamate**.

A suspension of 2-(aminomethyl)imidazo[1,2-*a*]pyridine-6-carbonitrile (4.2 g, 24.41 mmol) and *N*-ethyl-*N*-isopropyl-propan-2-amine (3.7 mL, 26.0 mmol) in THF (35 mL) and DCM (35 mL) was treated with *N*,*N*-dimethylpyridin-4-amine (0.085 g, 0.697 mmol) and *tert*-butoxy carbonyl *tert*-butyl carbonate (3.40 g, 15.6 mmol) at 0 °C and the mixture was allowed to warm to room temperature for 10 hours. The reaction mixture was concentrated under vacuum, the residue was suspended in water- $CH_3CN$ (1:1, 15mL), and sonicated for 20 min. The solid was filtered, and then treated with water-MeCN again. The obtained solid was dried under vacuum to give the title compound as an off white solid, which was further purify by the column chromatography on silica gel using eluent ethyl acetate in hexane to obtain *tert*-Butyl ((6-cyanoimidazo[1,2-a]pyridin-2-yl)methyl)carbamate (5.1 g, 86% yield).

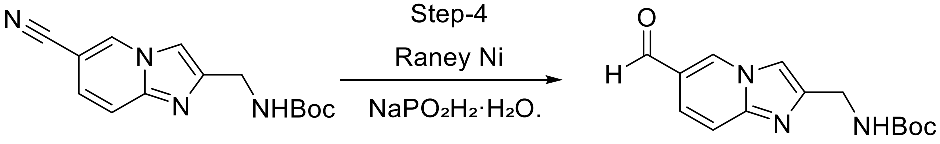


**Step 4: *tert*-Butyl ((6-formylimidazo[1,2-*a*]pyridin-2-yl)methyl)carbamate.**

A suspension of *tert*-butyl *N*-[(6-cyanoimidazo[1,2-a]pyridin-2- yl)methyl]carbamate (5.00 g, 18.4 mmol) and sodium phosphinate hydrate (15.57 g, 0.147 mol) in a mixture of water (50 mL), Pyridine (100 mL) and Acetic acid (50 mL) was treated with Raney nickel (50%, 18.97 g, 0.162 mol) and the mixture was heated at 100 °C under a nitrogen atmosphere for an hour. The Raney nickel was removed by hot filtration through a bed of Celite (washing with water, followed by methanol). The filtrate was concentrated under vacuum to remove the methanol and the blue solution was extracted with DCM (50 mL x 4). The extracts evaporated under vacuum to afford a beige gum which was triturated with water to furnish a white solid. The solid was collected by filtration, washed with water followed by ether and dried under vacuum overnight to provide the title compound *tert*-Butyl ((6-formylimidazo[1,2-a]pyridin-2-yl)methyl)carbamate (4.13 g, 82%) as an off white solid. $^1$H NMR (600 MHz, DMSO-$d_6$) $\delta$ 9.27 (s, 1H), 7.78 (s, 1H), 7.59 (dt, $J$ = 9.3, 0.9 Hz, 1H), 7.42 (dd, $J$ = 9.3, 1.7 Hz, 1H), 7.38 (t, $J$ = 6.1 Hz, 1H), 6.09 (s, 1H), 4.23 (d, $J$ = 6.0 Hz, 2H), 1.37 (s, 10H).

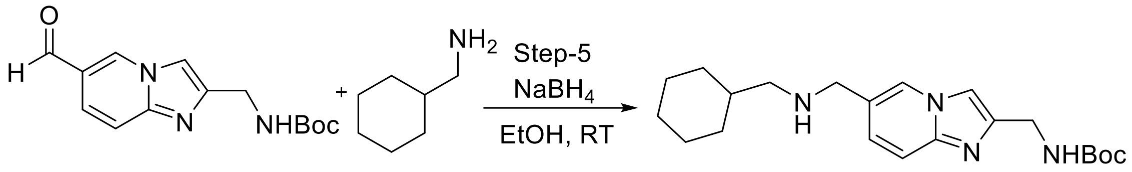


**Step 5: *tert*-Butyl ((6-(((cyclohexylmethyl)amino)methyl)imidazo[1,2-*a*]pyridin-2-yl)methyl)carbamate.**

A solution of *tert*-butyl *N*-[(6-formylimidazo[1,2-a]pyridin-2-yl)methyl]carbamate (340 mg, 0.988 mmol) and cyclohexyl methanamine (224 mg, 1.98 mmol) in ethanol (6.8 mL) was stirred at 50 °C for an hour. The reaction mixture was cooled to 0 °C and $NaBH_4$ (75 mg, 1.98 mmol) was added in portions. The reaction mixture was allowed to warm to room temperature and stirring was continued for 2 hours. The reaction mixture was partitioned between DCM (80 ml) and saturated aqueous sodium hydrogen carbonate (30 ml). The organic layer was separated, washed with brine (30 ml), dried over $MgSO_4$, concentrated under reduced pressure. The residue was purified by column chromatography on silica gel with 0-20% methanol in DCM to afford the title compound *tert*-Butyl ((6-(((cyclohexylmethyl)amino)methyl)imidazo[1,2-a]pyridin-2-yl)methyl)carbamate (360 mg, 97%) as a yellow oil. $^1$H NMR (600 MHz, DMSO-$d_6$) $\delta$ 8.37 (s, 1H), 7.65 (s, 1H), 7.39 (d, $J$ = 9.2 Hz, 1H), 7.27 (t, $J$ = 6.2 Hz, 1H), 7.19 (dd, $J$ = 9.2, 1.7 Hz, 1H), 4.21 (d, $J$ = 6.0 Hz, 2H), 3.63 (d, $J$ = 1.0 Hz, 2H), 2.31 (d, $J$ = 6.6 Hz, 2H), 1.75 – 1.71 (m, 2H), 1.66 – 1.59 (m, 3H), 1.40 (s, 9H), 1.37-1.34 (m, 1H), 1.26 – 1.01 (m, 4H), 0.87– 0.80 (m, 2H).

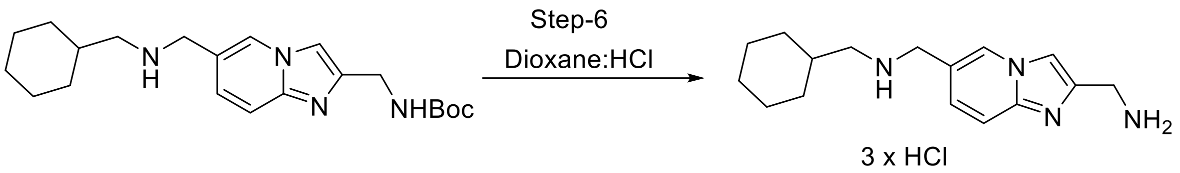


**Step 6. 1-(2-(Aminomethyl)imidazo[1,2-*a*]pyridin-6-yl)-*N*-(cyclohexylmethyl)methanamine**

To a solution of tert-butyl *N*-[[6-[(cyclohexylmethylamino)methyl]imidazo[1,2-a]pyridin-2-yl]methyl]carbamate (360 mg, 0.908 mmol) in methanol (4.1379 mL) was added HCl (4M in dioxane , 2.4 mL, 9.6 mmol) at room temperature The solution was stirred at 50 °C for 1 hour. The solution was cooled to room temperature, and concentrated to dryness under reduced pressure. The residue (pale yellow glass) was dissolved in methanol (~3 mL) and diethyl ether (~ 20 mL) was added drop-wise to the stirred solution. The resulting precipitate was collected by filtration and dried in the vacuum oven at 40 °C for 4 hours to afford the title compound 1-(2-(Aminomethyl)imidazo[1,2-a]pyridin-6-yl)-*N*-(cyclohexylmethyl)methanamine trihydrochloride salt (339 mg, 95%) as white solid.

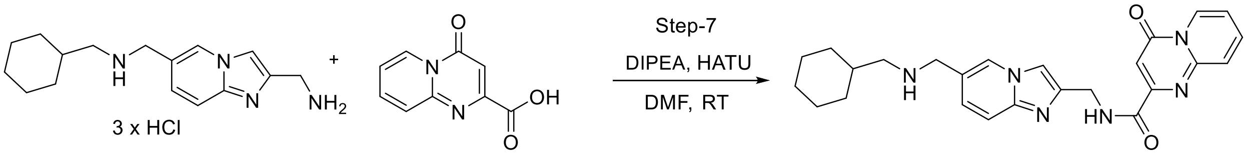


**Step 7. *N*-((6-(((Cyclohexylmethyl)amino)methyl)imidazo[1,2-*a*]pyridin-2-yl)methyl)-2-oxo-2*H*-pyrido[1,2-*a*]pyrimidine-4-carboxamide**

A solution of 4-oxopyrido[1,2-a]pyrimidine-2-carboxylic acid (242 mg, 1.2 mmol), HATU (724 mg, 1.9 mmol) and 1-(2-(aminomethyl)imidazo[1,2-a]pyridin-6-yl)-*N*-(cyclohexylmethyl) methanamine trihydrochloride (414 mg, 1.5 mmol) in DMF (7 mL) was treated with *N*-ethyl-*N*-isopropyl-propan-2-amine (1.1 mL, 8 mmol) and the suspension was stirred at room temperature for 16 hours. The suspension was concentrated under vacuum and the residue was diluted with water. The solids were then collected by filtration, washed with water followed by ether and dried under vacuum to give the title compound (203 mg, 36%) as an off-white solid. $^{1}$H NMR (600 MHz, DMSO-$D_6$) $\delta$ 9.14 (t, $J$ = 6.0 Hz, 1H), 9.02 (d, $J$ = 6.5 Hz, 1H), 8.37 (s, 1H), 8.07 (ddd, $J$ = 8.6, 6.7, 1.6 Hz, 1H), 7.80 (dt, $J$ = 8.9, 1.1 Hz, 1H), 7.78 – 7.76 (m, 1H), 7.51 – 7.41 (m, 2H), 7.21 (dd, $J$ = 9.3, 1.7 Hz, 1H), 6.90 (s, 1H), 4.61 (d, $J$ = 5.9 Hz, 2H), 3.64 (s, 2H), 2.30 (d, $J$ = 6.6 Hz, 2H), 1.76 – 1.70 (m, 2H), 1.66 – 1.58 (m, 3H), 1.43 – 1.34 (m, 1H), 1.19 – 1.08 (m, 3H), 0.86 – 0.81 (m, 2H). $^{13}$C NMR (150 MHz, DMSO-$D_6$) $\delta$ 162.66, 157.85, 153.68, 150.45, 143.70, 143.39,

138.42, 127.36, 126.18, 126.03, 124.92, 124.34, 117.21, 115.75, 110.08, 100.72, 55.12, 50.13, 37.67, 37.50, 31.01, 26.25, 25.58.

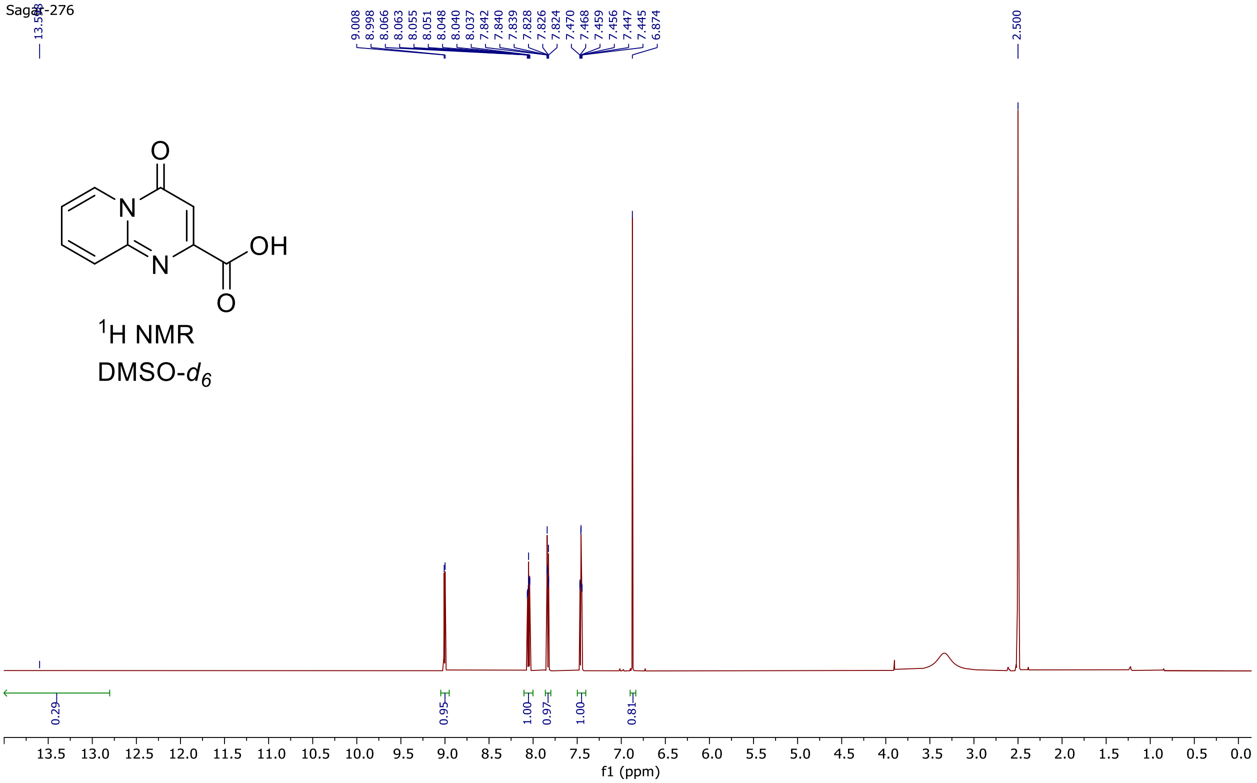


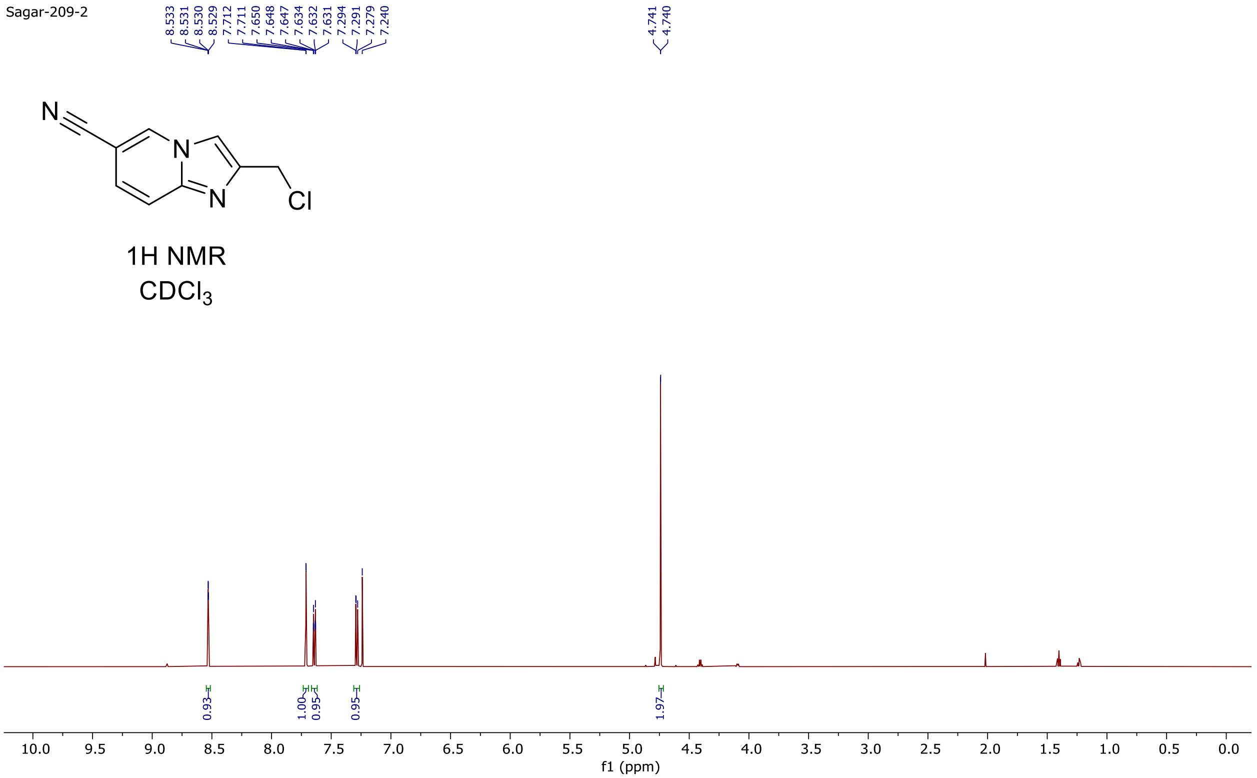

Sagar-217

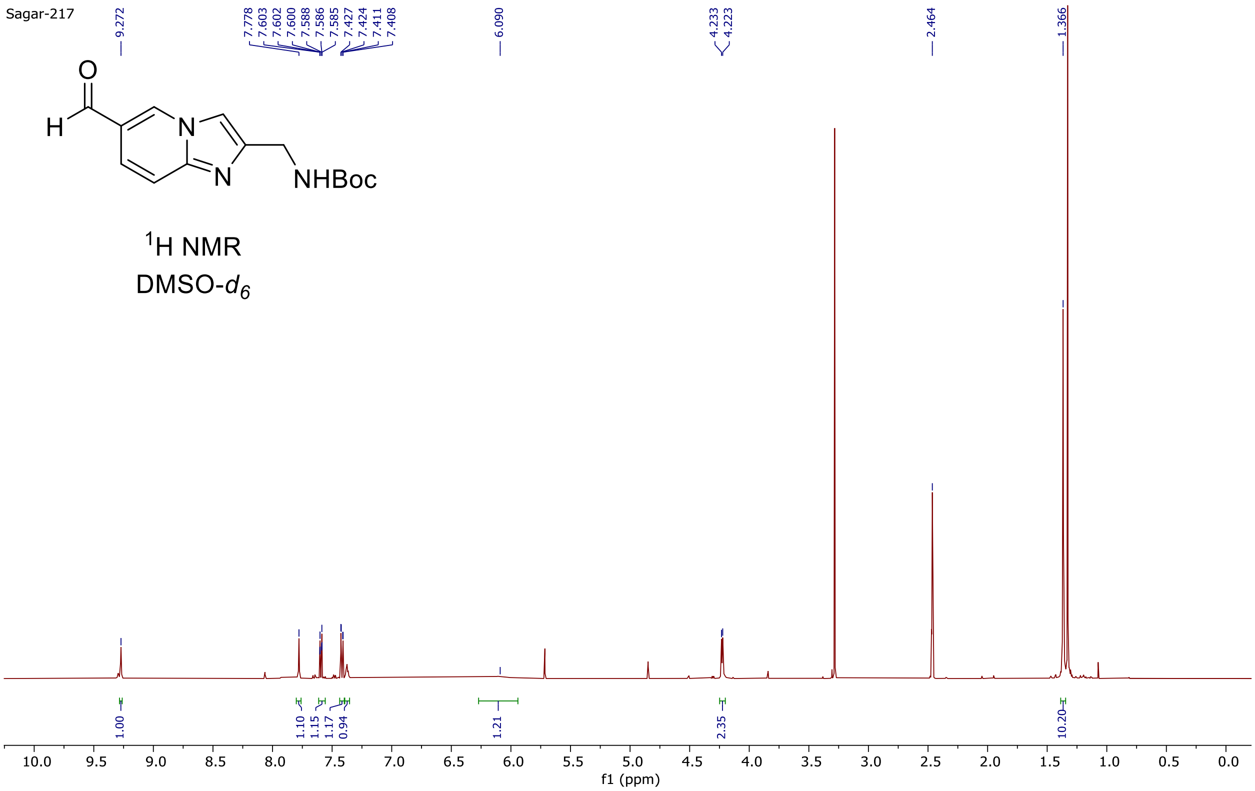


$^1$H NMR
DMSO-$d_6$

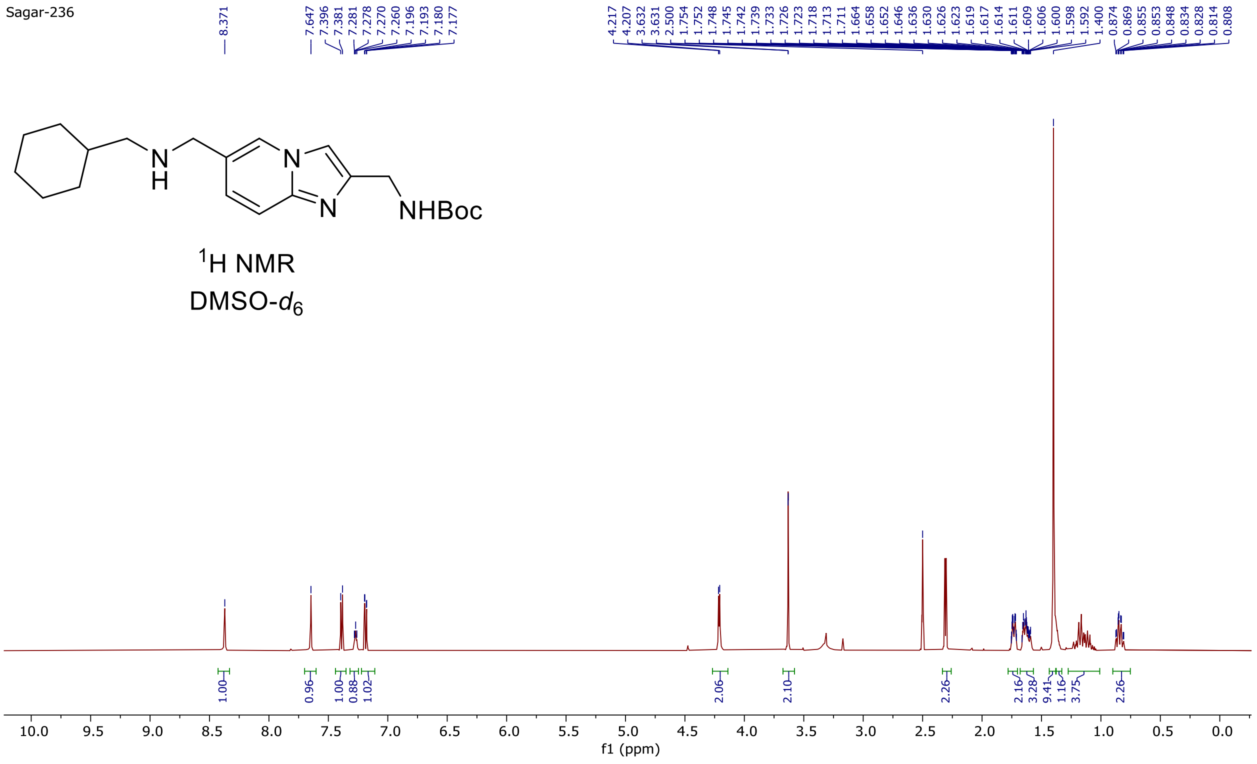
Sagar-236
8.371
7.647 7.396 7.381 7.281 7.278 7.270 7.260 7.196 7.193 7.180 7.177
4.217 4.207 3.632 3.631 2.500 1.754 1.752 1.748 1.745 1.742 1.739 1.733 1.726 1.723 1.718 1.713 1.711 1.664 1.658 1.652 1.646 1.636 1.630 1.626 1.623 1.619 1.617 1.614 1.611 1.609 1.606 1.600 1.598 1.592 1.400 0.874 0.869 0.855 0.853 0.848 0.834 0.828 0.814 0.808
N
H
N
N
NHBoc
1H NMR
DMSO-d6
1.00 0.96 1.00 0.88 1.02 2.06 2.10 2.26 2.16 3.28 9.41 1.16 3.75 2.26
10.0 9.5 9.0 8.5 8.0 7.5 7.0 6.5 6.0 5.5 5.0 4.5 4.0 3.5 3.0 2.5 2.0 1.5 1.0 0.5 0.0
f1 (ppm)

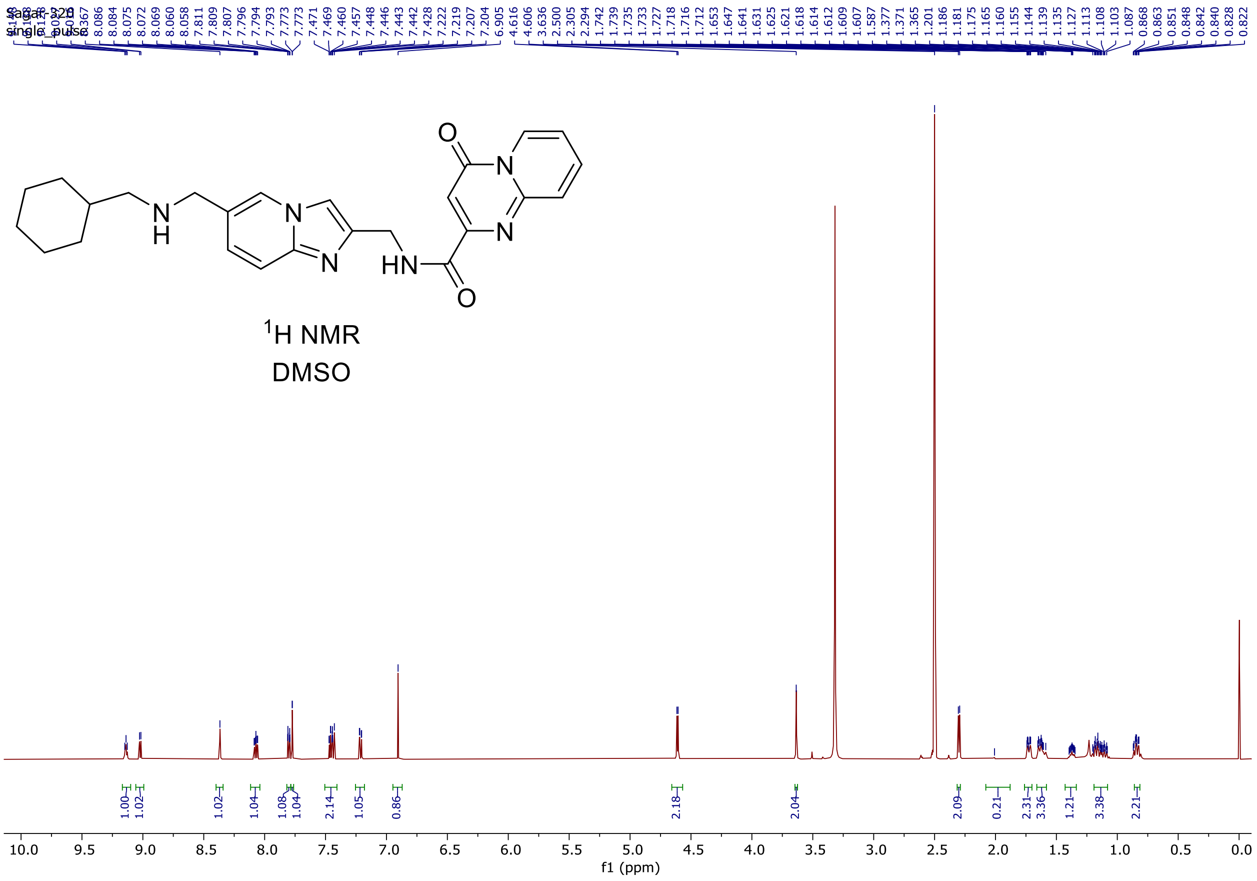
Sagar-320
single_pulse
8.367 8.086 8.084 8.075 8.072 8.069 8.060 8.058 7.811 7.809 7.807 7.796 7.794 7.793 7.773 7.773 7.471 7.469 7.460 7.457 7.448 7.446 7.443 7.442 7.428 7.222 7.219 7.207 7.204 6.905 4.616 4.606 3.636 2.500 2.305 2.294 1.742 1.739 1.735 1.733 1.727 1.716 1.712 1.653 1.647 1.641 1.631 1.625 1.621 1.618 1.614 1.612 1.609 1.607 1.587 1.377 1.371 1.365 1.201 1.186 1.181 1.175 1.165 1.160 1.155 1.144 1.139 1.135 1.127 1.113 1.108 1.103 1.087 0.868 0.863 0.851 0.848 0.842 0.840 0.828 0.822
O
N
N
N
H
N
N
HN
O
1H NMR
DMSO
1.00 1.02 1.02 1.04 1.08 1.04 2.14 1.05 0.86 2.18 2.04 2.09 0.21 2.31 3.36 1.21 3.38 2.21
10.0 9.5 9.0 8.5 8.0 7.5 7.0 6.5 6.0 5.5 5.0 4.5 4.0 3.5 3.0 2.5 2.0 1.5 1.0 0.5 0.0
f1 (ppm)

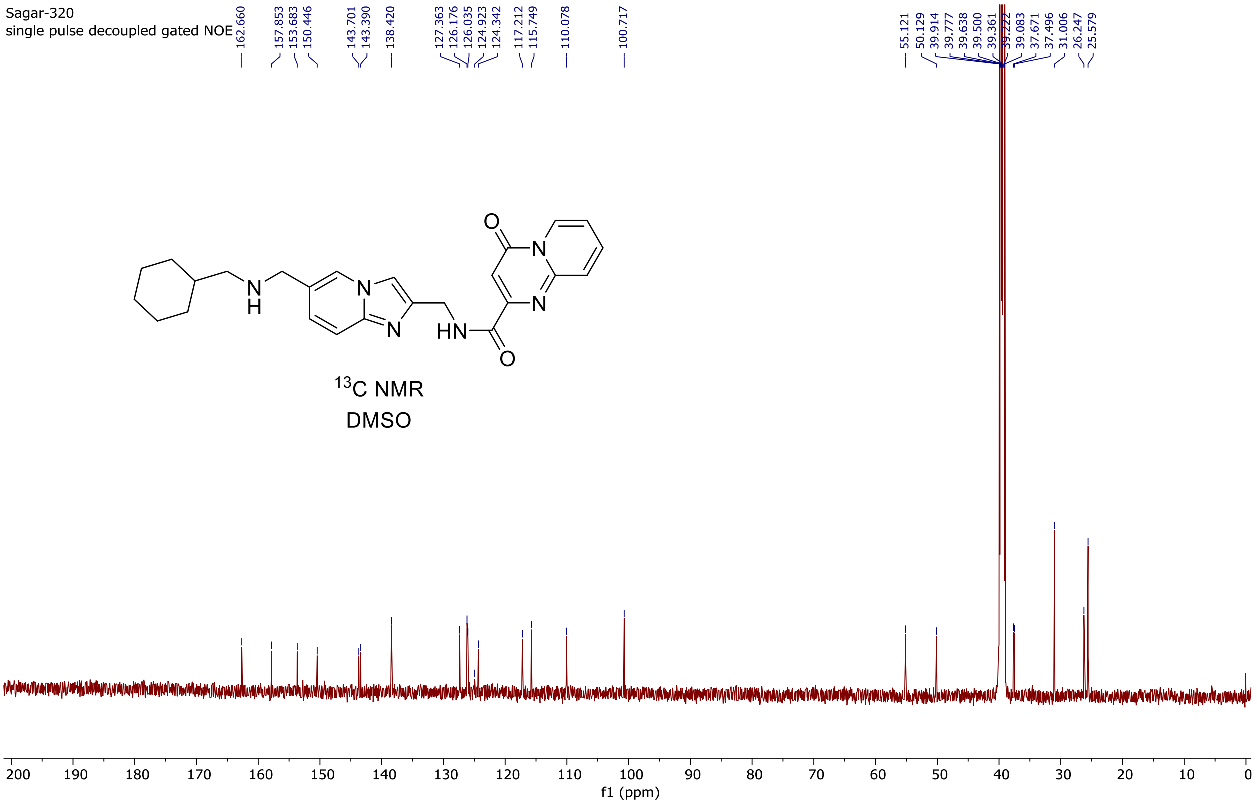

Sagar-320
single pulse decoupled gated NOE
162.660
157.853
153.683
150.446
143.701
143.390
138.420
127.363
126.176
126.035
124.923
124.342
117.212
115.749
110.078
100.717
55.121
50.129
39.914
39.777
39.638
39.500
39.361
39.222
39.083
37.671
37.496
31.006
26.247
25.579
13C NMR
DMSO
f1 (ppm)
